# Inhibiting adult neurogenesis differentially affects spatial learning in females and males

**DOI:** 10.1101/2021.10.27.466135

**Authors:** Timothy P O’Leary, Baran Askari, Bonnie Lee, Kathryn Darby, Cypress Knudson, Alyssa M Ash, Desiree R Seib, Delane F Espinueva, Jason S Snyder

**Affiliations:** Department of Psychology, Djavad Mowafaghian Centre for Brain Health, University of British Columbia, Vancouver, BC, Canada

## Abstract

Adult hippocampal neurogenesis has been implicated in the spatial processing functions of the hippocampus but ablating neurogenesis does not consistently lead to behavioral deficits in spatial tasks. Parallel studies have shown that adult-born neurons also regulate behavioral responses to stressful and aversive stimuli. We therefore hypothesized that spatial functions of adult-born neurons may be more prominent under conditions of stress, and may differ between males and females given established sex differences in stress responding. To test this we trained intact and neurogenesis-deficient rats in the spatial water maze at temperatures that vary in their degree of aversiveness. At standard temperatures (25°C) ablating neurogenesis did not alter learning and memory in either sex, consistent with prior work. However, in cold water (16°C), ablating neurogenesis had divergent sex-dependent effects: relative to intact rats, male neurogenesis-deficient rats were slower to escape and female neurogenesis-deficient rats were faster. Neurogenesis promoted temperature-related changes in search strategy in females, but it promoted search strategy stability in males. Females displayed greater recruitment of the dorsal hippocampus than males, particularly at 16°C. However, blocking neurogenesis did not alter activity-dependent immediate-early gene expression in either sex. Finally, morphological analyses of retrovirally-labelled neurons revealed greater experience-dependent plasticity in new neurons in males. Neurons had comparable morphology in untrained rats but 16°C training increased spine density, and 25°C training caused shrinkage of mossy fiber presynaptic terminals, specifically in males. Collectively, these findings indicate that neurogenesis functions in memory are prominent under conditions of stress, they provide the first evidence for sex differences in the behavioral function of newborn neurons, and they suggest possibly distinct roles for neurogenesis in cognition and mental health in males and females.

## INTRODUCTION

Adult hippocampal neurogenesis has been implicated in many of the mnemonic functions of the hippocampus, including memory for temporal events (1–3), locations (4), contexts (5, 6), objects (7, 8) and conspecifics (9), as well as the consolidation (10, 11) and forgetting (12) of memory. While spatial memory functions may be particularly apparent in conditions that maximize conflict or interference, such as when a goal changes location (13–16), it is notable that many of studies have failed to find a role for new neurons in learning and short-term reference memory in the spatial water maze, a task that is highly sensitive to hippocampal disruption (2, 6, 7, 17–22).

A relatively independent body of work has focused on the role of neurogenesis in emotional and stress-related behavior, finding that neurogenesis buffers the endocrine response to acute stressors and reduces depressive- and anxiety-like behavior (23–29). Since stress and emotion potently modulate learning and memory (30, 31), here we hypothesized that a role for neurogenesis in spatial learning may become particularly apparent in more aversive conditions. Consistent with this possibility, a small number of studies have found that neurogenesis does alter behavior in memory tasks depending on the aversiveness of conditioned and unconditioned stimuli that are present (3, 32, 33).

Stress-related disorders such as anxiety, PTSD and depression impact a substantial fraction of the population. Critically, these disorders affect females to a greater extent than males, suggesting that neurogenesis functions in stress may be particularly relevant for female cognition and mental health (34). Indeed, there are known sex differences in the rates of addition (35), maturation (36) and activation of adult-born neurons (37). Furthermore, sex modulates hippocampal plasticity (38–41) and behavioral responses to acute and chronic stress (42–44). However, as is the case in neuroscience more broadly (45), the majority of neurogenesis studies have focused on males (46). In a quantitative survey of the neurogenesis literature we find that males are studied twice as often as females, less than 10% of studies have reported data by sex, and more than 20% of studies do not report the sex of their subjects (Fig. 1). To our knowledge, no sex differences in behavior have been reported in animals that have specific reductions in adult neurogenesis. One study has found reduced neurogenesis is associated with female-specific impairments in adult learning (47). However, this was in response to neurogenesis ablation beginning in infancy, raising the question of whether sex differences may also occur in response to neurogenesis ablation in adulthood.

**Figure 1:**
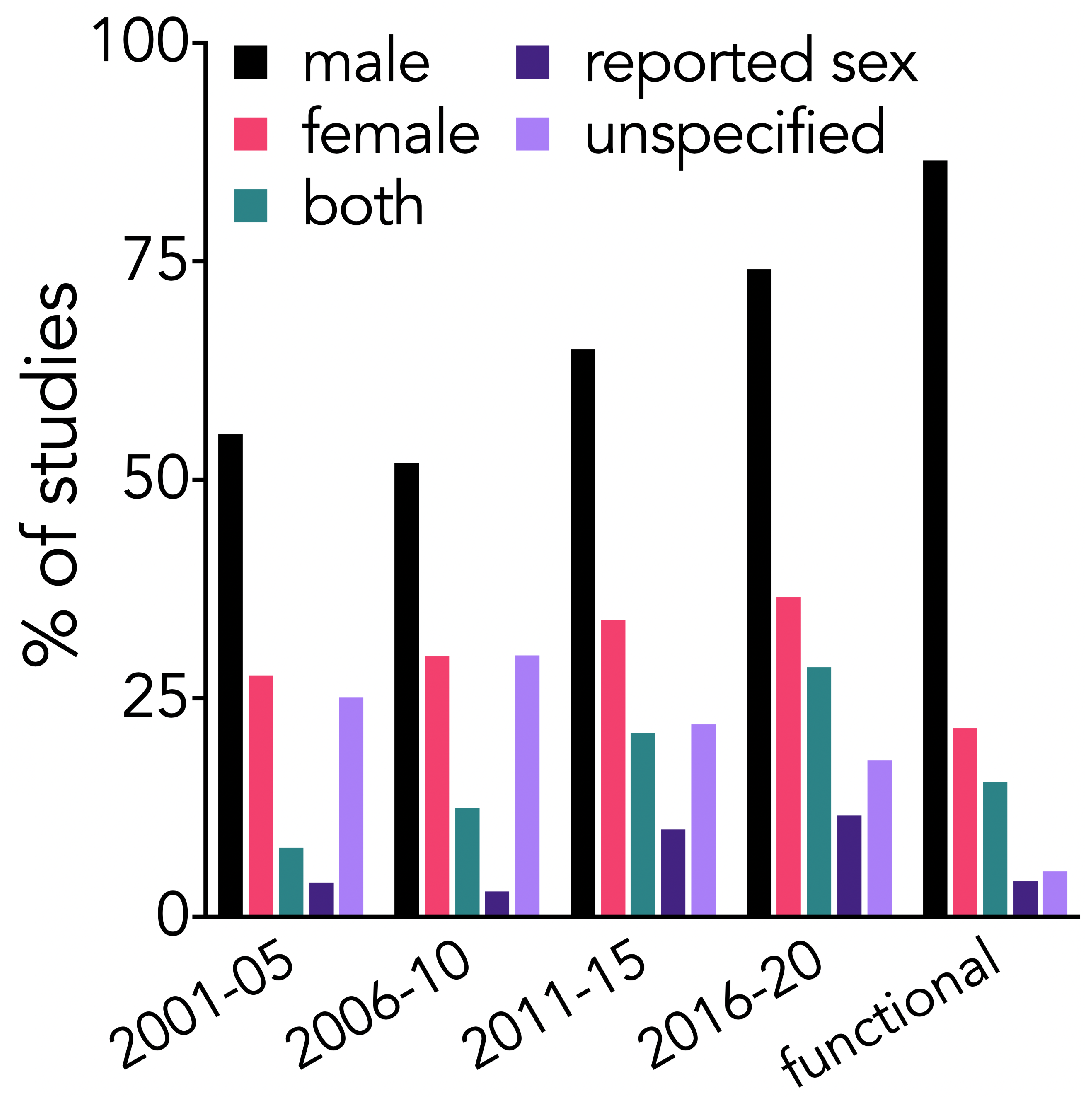
Neurogenesis studies primarily include males. Over the past 20 years, most studies have included males, fewer have included females or specifically reported data by sex. A fraction continue to not specify the sex of their subjects. Similar patterns are seen for “functional” studies that have manipulated neurogenesis and examined behavioral outcomes.

To address these outstanding issues we used a pharmacogenetic GFAP-TK rat model to block adult neurogenesis (48), and tested male and female rats in the water maze at warm (25°C, standard) or cold (16°C, more aversive/stressful) temperatures. Consistent with previous work, neurogenesis-deficient rats were unimpaired at standard water maze temperatures. However, cold water testing revealed striking sex differences in the behavioral function of adult neurogenesis, and also elicited distinct dorsoventral patterns of hippocampal recruitment and new neuron plasticity in males and females.

## METHODS

### Subjects

This study used male and female transgenic GFAP-TK (“TK”) and wild-type (“WT”) rats on a Long-Evans background (48). Here, a GFAP promoter drives expression of herpes simplex virus thymidine kinase in radial-glial precursor cells, enabling these cells to be killed when rats are treated with valganciclovir and the cells attempt mitosis. Rats were bred in-house, by crossing heterozygous transgenic females with WT males. After weaning (postnatal day 21) rats were housed in same-sex groups of 2-3 in polyurethane cages (48 cm x 27 cm x 20 cm), with aspen chip bedding, a polycarbonate tube for enrichment, and ad-libitum access to food and water. Animals were housed under a 12-hour light:dark cycle, and all testing was completed during the light phase. Rats were genotyped via PCR after weaning and, therefore, housed randomly with respect to genotype. Prior to all experiments, animals were handled 5 min/day for 5 days. Experimental procedures were approved by the University of British Columbia Animal Care Committee and followed guidelines from the Canadian Council of Animal Care on the ethical treatment of animals.

### Valganciclovir treatment

For experiments with neurogenesis ablation, animals were treated orally with pellets of valganciclovir (4 mg) in a 1:1 peanut butter and rodent chow mix (0.5 g). Drug pellets were given directly to each animal to ensure accurate dosing. Animals began treatment at 6-7 weeks of age, and were treated twice a week (3-4 day interval) for 6-7 weeks before behavioral testing began. Valganciclovir treatment stopped immediately prior to behavioral testing. In control experiments without neurogenesis ablation, rats received neither valganciclovir nor peanut butter and rodent chow mix.

### Spatial water maze testing

The water maze consisted of a white circular pool (180 cm diameter), with 60 cm high walls. The pool was filled with water to a 32 cm depth, and the water was made opaque with addition of white non-toxic liquid tempera paint (Schola). Training contexts of high- or moderate-stress were created by using either 16°C or 25°C water, respectively, similar to previous work (49, 50). The pool was located in a room (~4m x 6m in size) with diffuse lighting, and contained extramaze visual cues along the room’s walls and distributed within the room (desk, computer, cabinets). A circular escape platform (12 cm diameter) was placed in the NE quadrant of the pool, and was positioned 2 cm below the water surface. Rats received 3 days of acquisition training with 4 trials per day. Rats were tested in groups of 2-3, and during daily training sessions were placed into individual holding cages filled with aspen chip bedding and paper towel.

For each trial, rats were placed into the pool at one of four possible release locations (pseudo-random order), with each release location occurring once on each day. Rats were given a maximum of 60 sec to locate the escape platform, after which they were guided to the escape platform by the experimenter. Following each trial, rats remained on the escape platform for ~10 sec, and were gently dried with a towel before being returned to their holding cage for the inter-trial interval (30-90 sec). The rats’ trajectory was recorded with an Ethovision (Noldus) tracking system, and performance was assessed via latency to locate the escape platform and swim speed. Ideal path error (conceptually similar to cumulative search error / proximity metrics (51)), which can detect spatial performance differences between trials that have similar latencies and distances (52), was calculated with Pathfinder software as follows: the distance from the platform was summed over all samples to obtain a cumulative distance metric. To control for different release locations, the cumulative distance for the optimal path was also calculated based on a direct escape path from the release location and the average swim speed. The ideal path error was then calculated by subtracting the cumulative optimal path from the cumulative actual path. On the day following acquisition training the platform was removed from the pool and rats completed a 60 sec probe trial to assess memory. Spatial memory was measured as the time spent in a 36 cm zone surrounding the former escape platform location, and the corresponding 36 cm zones in each of the non-target quadrants. Rats were euthanized 60 minutes after the probe trial in order to capture experience-dependent Fos expression in activated neurons (see *Immunohistochemistry*, below).

### Search strategy analyses

Navigational search strategies employed in the water maze were detected using Pathfinder software (52), with the following parameters: angular corridor width: 45°, chaining annulus width: 45cm, thigmotaxis zone width: 15cm, Direct Swim maximum ideal path error: 125, max heading error: 35°; Focal Search max distance to swim path centroid: 30, max distance to goal: 30, min distance covered: 100cm, max distance covered: 500cm; Directed Search min time in angular corridor: 70%, max distance covered: 400cm, max ideal path error: 1500; Indirect Search max ideal path error: 450, max average heading error: 70°; Semi-Focal Search max distance to swim path centroid: 60, max distance to goal: 60, min distance covered: 200cm, max distance covered: 5000cm; Chaining min time in annulus: 70%, min quadrants visited: 4, max area of maze traversed: 40%; Scanning max area of maze traversed: 20%, min area of maze traversed: 0%, max average distance to maze center: 60; Thigmotaxis time in full zone: 60%, time in smaller zone: 0%, min total distance covered: 400cm. Random search min area of maze traversed: 5%. The small number of trials that were not categorized by Pathfinder were designated as random.

To visualize the change in usage of a given strategy, S, caused by reducing neurogenesis, weighted difference scores were calculated as: (% trials S_TK_ - % trials S_WT_)/% trials S_WT_) × (# trials S_WT+TK_)/(# trials total_WT+TK_) × 100. In other words, strategy difference scores were weighted against their relative frequency, to prevent overrepresentation of differences that occurred on only a small proportion of the total trials.

### Retrovirus injections

Moloney Murine Leukemia-Virus retrovirus, produced as recently described (53), was use to express eGFP in adult-born neurons. Viral titers ranged from 1 to 8 x 10^6^ colony forming units/ml. Eight-week-old male and female rats were bilaterally injected with 1 μl of retrovirus into the dorsal dentate gyrus (anteroposterior = −4.0 mm; mediolateral = ±3.0 mm; dorsoventral = −3.5 mm from bregma). Thirty days later, rats either remained in their home cage or were trained and tested for 4 days in the 16°C or 25°C water maze, as above. Rats were perfused the next day, when cells were 35 days old.

### Blood sampling and radioimmunoassays

In one group of rats, different from those used to generate the main behavioral data in Figures 2–3, blood samples were obtained 30 min following testing sessions on days 1 and 3 of acquisition training and after the probe trial on day 4. After the last trial of a training session was completed, rats remained in the testing room for 5 min, before being returned to their home-cage and colony room for the remaining 25 min. Rats were then quickly brought into the hallway adjacent to the colony room, restrained, and blood was collected via tail vein puncture. For baseline circadian measurements, home cage control rats were sampled directly from their cage without transport. Blood was left at room temperature for 30-45 min, centrifuged, and serum supernatant was collected and stored at −80°C until analyzed by radioimmunoassay (RIA). RIAs were completed using a I^125^ corticosterone competitive binding assay (MP Biomedical). In a subset of animals, body temperature was also obtained immediately following blood sampling using a rectal thermometer.

**Figure 2:**
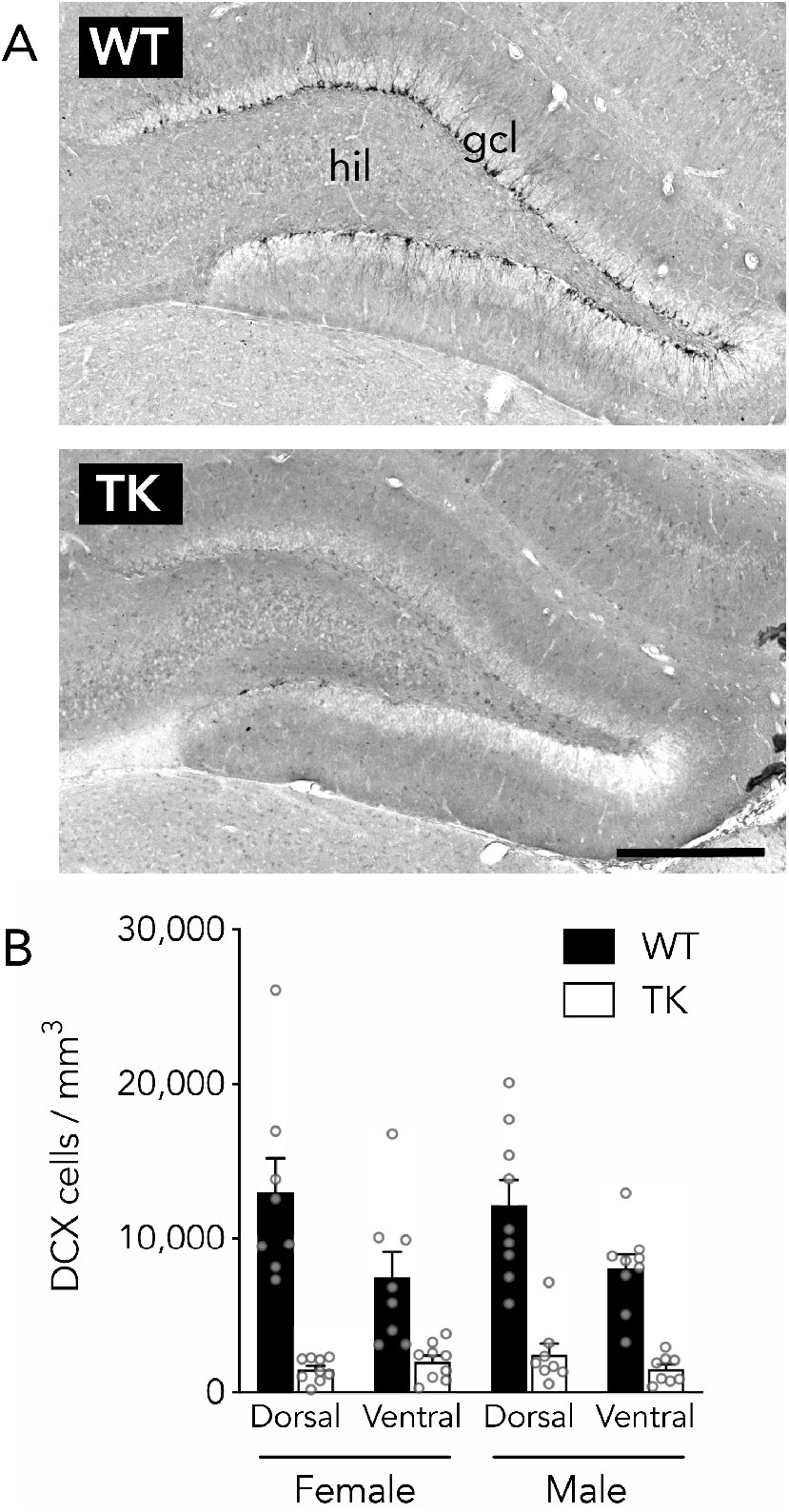
Reduced neurogenesis in male and female GFAP-TK rats. A) Representative immunostaining for the immature neuronal marker, doublecortin (DCX), in WT (top) and TK (bottom) rats (here, both female). hil, hilus; gcl, granule cell layer; scale bar, 500 μm. B) Neurogenesis was suppressed along the dorsoventral axis of both male and female rats (3 way ANOVA; effect of genotype: F_1,30_ = 58, P<0.0001; effect of sex: F_1,30_ = 0.0, P=0.96; effect of dorsoventral subregion: F_1,30_ = 28, P<0.0001; interactions all P>0.15). Bars reflect mean ± standard error.

**Figure 3:**
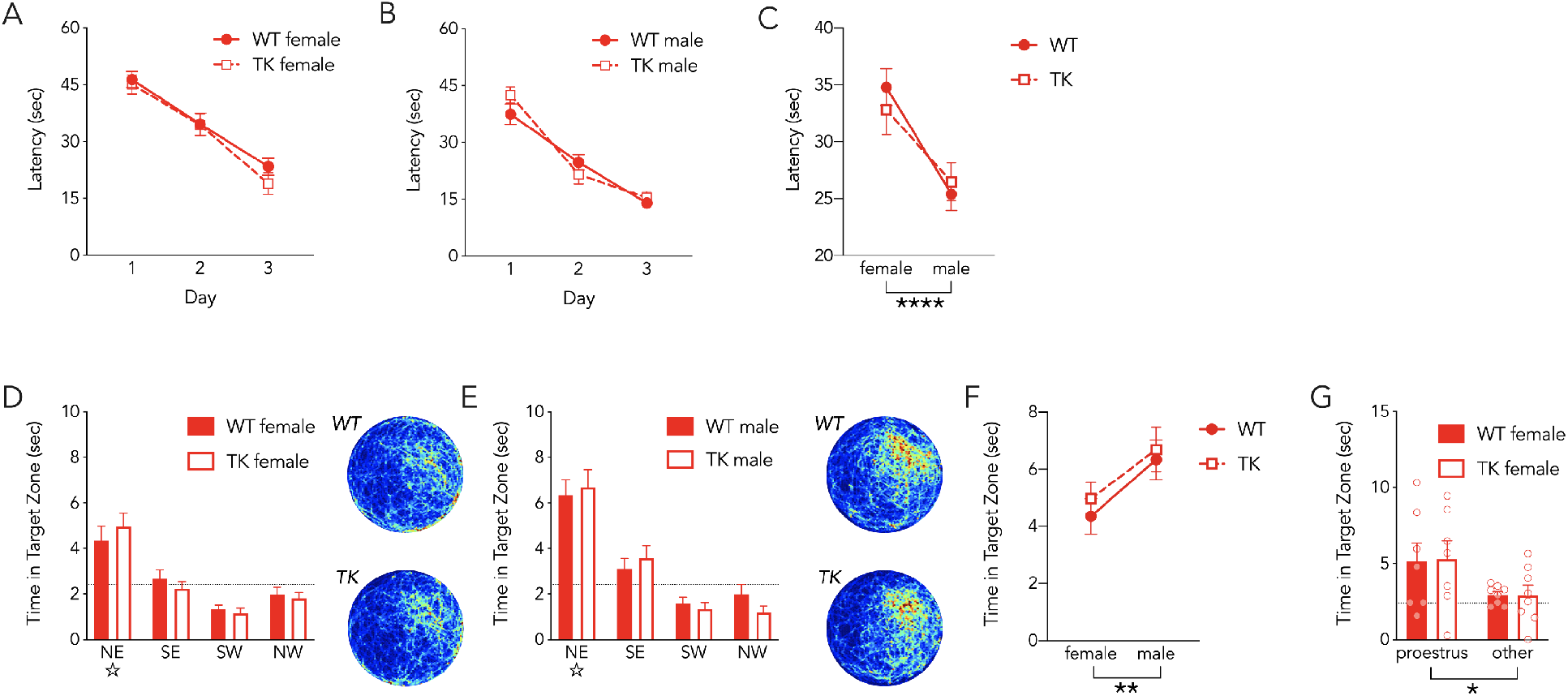
Sex, but not neurogenesis, modulates learning and memory in the 25°C water maze. During the acquisition phase of water maze training, female (A) and male (B) rats reached the platform faster with successive days of training (3-way anova; effect of day, F_2,152_=157, P<0.0001). Males reached the platform faster than females (effect of sex, F_1,76_=21, P<0.0001) but there was no difference between WT and TK rats (effect of genotype, F_1,76_=0.1, P=0.8) and no significant interactions between day, sex and genotype (all P > 0.1). C) Summary of average trial acquisition latency. D, E) On the probe trial, female (D) and male (E) rats preferentially searched in the target (NE) zone where the platform was located during training. Dotted line indicates chance performance. WT and TK rats did not differ on the probe trial but males spent more time searching in the target zone (2-way anova; effect of genotype, F_1,69_ = 0.4, P=0.5; effect of sex, F_1,69_=7, P=0.0097; interaction, F_1,69_=0, P=0.8). F) Summary of probe trial target zone search time for males and females. G) Females in proestrus displayed better memory on the probe trial than rats in other phases of the estrous cycle (2-way anova; effect of genotype, F_1,26_ = 0, P=0.9; effect of estrous stage, F_1,26_= 6.5, P=0.02; interaction, F_1,26_=0, P=0.8). **P<0.01, ****P<0.0001. N=18-22 per group. Bars and symbols reflect mean ± standard error. Heat maps scaled equivalently for males and females.

### Vaginal lavage and estrous staging

Vaginal lavages were completed on female rats within 1-6 hours of completing the probe trial. Rats were gently wrapped in a towel and rotated so that the vagina was clearly visible. The vagina was then flushed with tap water using a glass transfer pipette with a smooth, curved tip. The water was then aspirated into the pipette and collected on a glass slide. The samples were left to dry for at least 24 hours before being stained in cresyl violet (0.1% for 1 min). For animals that were used in Figures 3–6, lavages were performed immediately prior to euthanasia and perfusion, to prevent any effects of lavage on water maze behavior or experience-dependent Fos expression. Additionally, only a portion of the animals that were used for these figures were lavaged. For animals that were used for corticosterone measurements, lavage was performed at the same time blood was collected. Identification of estrous cycle stage was completed based on the cytology of lavages, as described (54), using an Olympus CX41 light microscope. Briefly, proestrus was identified based on the presence of round squamish cells with visible nuclei, estrous with cornified squamish cells without visible nuclei, metestrus with both cornified squamish cells and leukocytes and diestrus with squamish cells that have visible nuclei and leukocytes.

**Figure 4:**
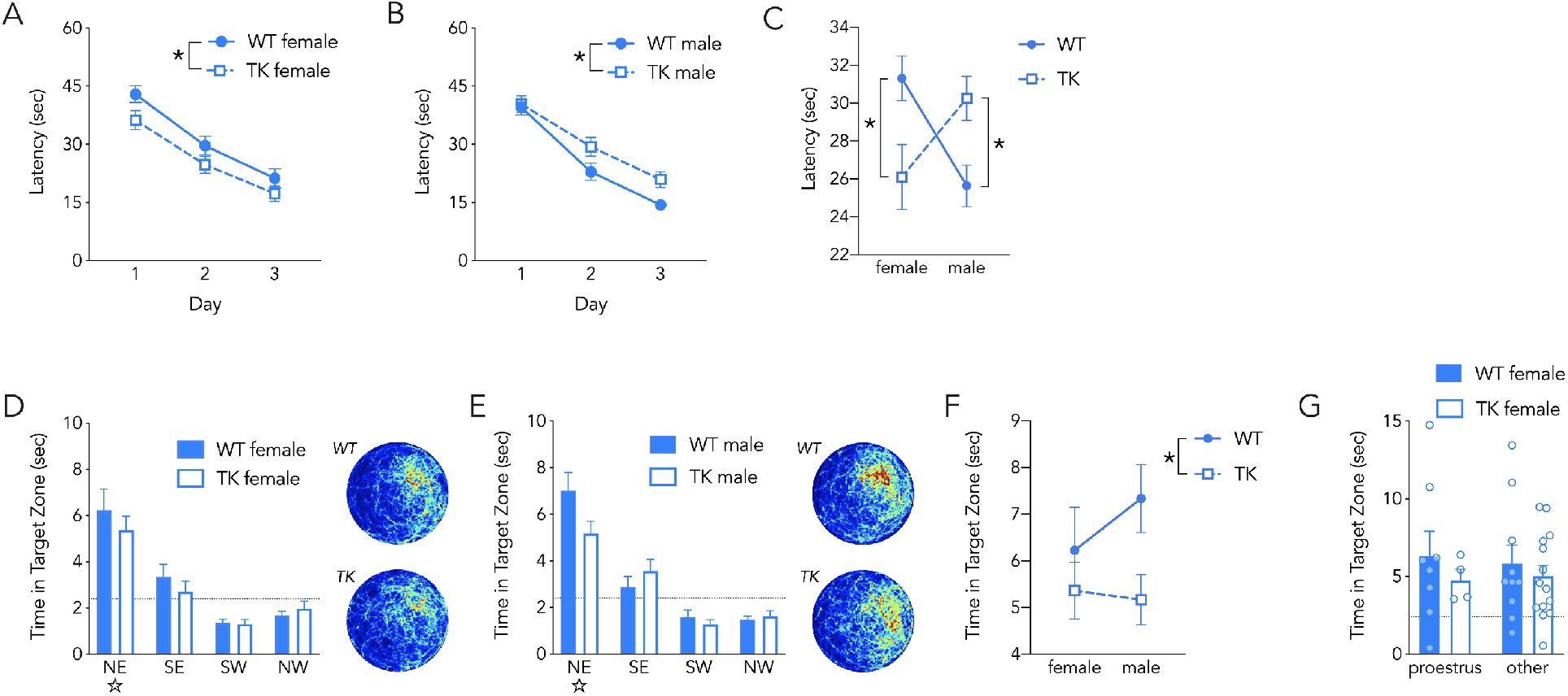
Sex and neurogenesis regulate learning and memory in the 16°C water maze. During acquisition, female (A) and male (B) reached the platform faster with successive days of training (3-way anova; effect of day, F_2,156_=105, P<0.0001). There was no main effect of sex (F_1,78_=0.3, P=0.6) or genotype (F_1,78_=0.1, P=0.8) but there was a significant sex x genotype interaction (F_1,78_=15, P=0.0003). Post hoc tests revealed that male WT rats learned faster than male TK rats (P=0.01) and female WT rats learned slower than female TK rats (P=0.01). C) Summary of average acquisition latencies in females and males. On the probe trial, female (D) and male (E) rats preferentially searched in the target (NE) zone where the platform was located during training. Dotted line indicates chance performance. There was no effect of sex on probe trial performance, but WT rats spent more time searching in the correct zone than TK rats (2-way anova; effect of genotype, F_1,76_=5, P=0.03; effect of sex, F_1,76_=0.4, P=0.4; interaction, F_1,76_ = 0.9, P=0.4). F) Summary of time spent in the target zone of the probe trial for females and males. G) Estrous stage did not influence performance on the probe trial (2-way anova; effect of genotype, F_1,32_=0, P=0.9; effect of estrous stage, F_1,32_=0, P=0.9; interaction, F_1,32_ =0, P=0.7). **P<0.01, ****P<0.0001. N=18-22 per group. Bars and symbols reflect mean ± standard error. Heat maps for females and males are scaled equivalently and match those in Fig 2.

**Figure 5:**
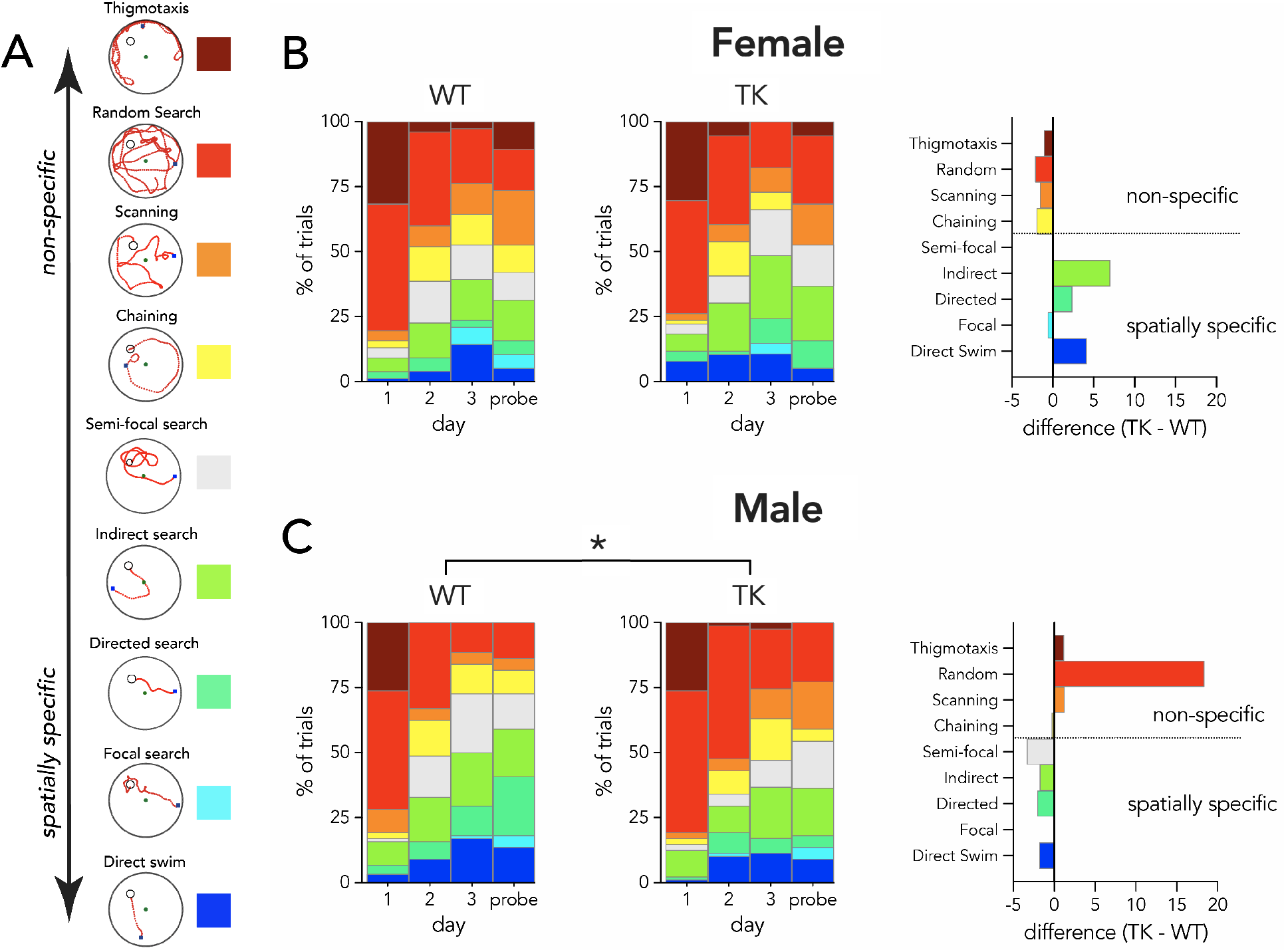
In the 16°C water maze, blocking neurogenesis reduces spatially-specific search in male rats. A) Example trials illustrating various search strategies classified by Pathfinder, organized by degree of spatial specificity relative to the target. B) Strategies employed by female WT and TK rats. The distribution of strategies in female TK rats was not significantly different from female WT rats (**χ^2^** = 7, P = 0.5). Right-most graph shows weighted strategy changes in TK rats relative to WT rats, quantified as % changes in strategy usage multiplied by the total fraction of trials where rats employed that strategy (genotypes pooled). The magnitude of the bars therefore reflects changes in strategy but prevents misleading perceptual artefacts caused by large % changes for strategies that were rarely used. (C) Reducing neurogenesis significantly altered the distribution of strategies used by male rats, demonstrated by the greater proportion of spatially non-specific trials and the smaller proportion of spatially-specific trials (right; **χ^2^** = 17, P = 0.02).

**Figure 6:**
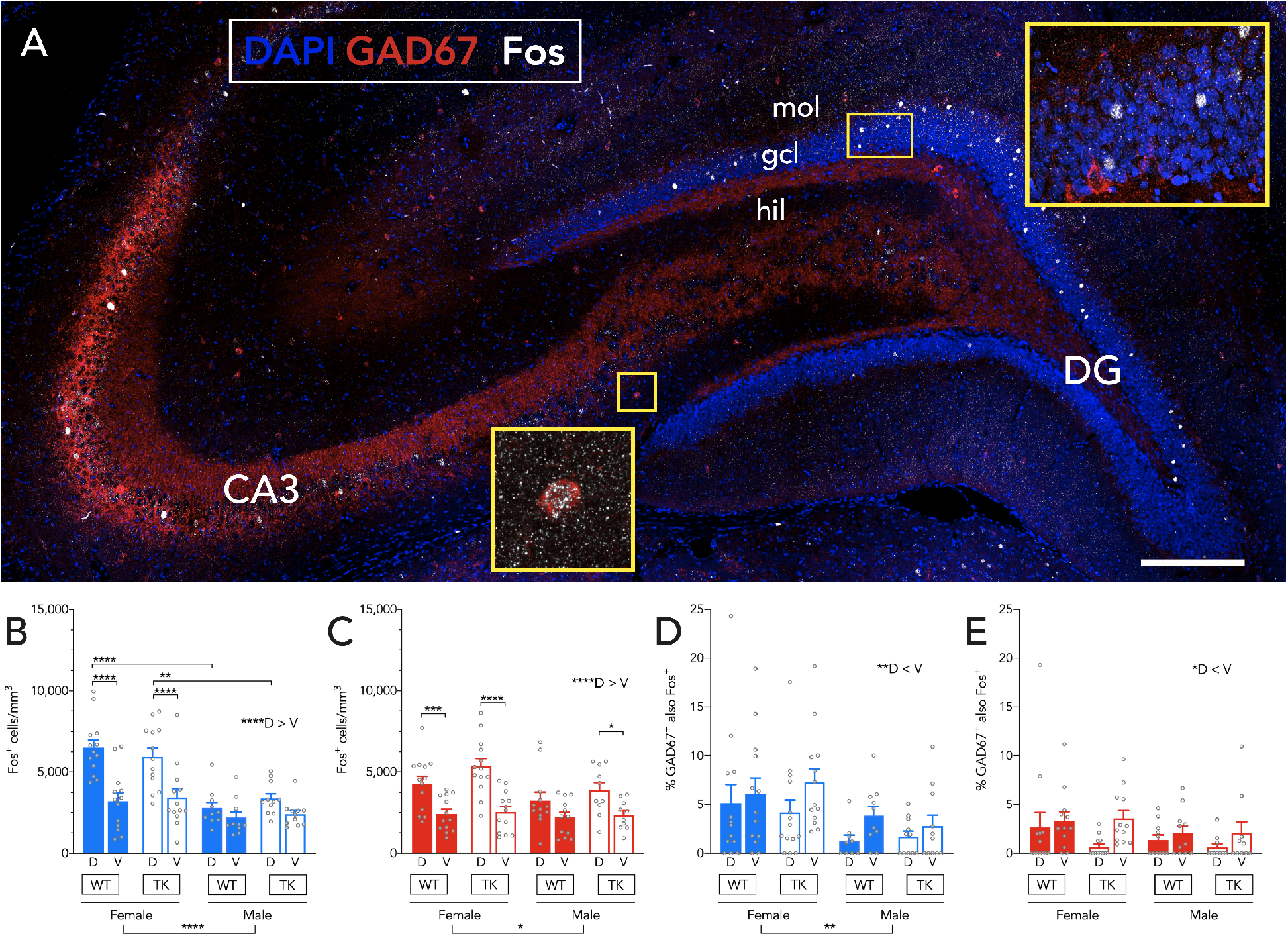
Sex- and subregion-based activation of DG-CA3 neurons. A) Confocal image of dorsal hippocampus immunostained for GAD67 and Fos. Scale bar, 200 μm. B) In the 16°C water maze, there were more Fos^+^ cells in the dorsal granule cell layer, particularly in female rats. The density of Fos^+^ cells was also greater in the dorsal DG of females that in males (3 way RM ANOVA; effect of subregion: F_1,43_=52, P<0.0001; effect of sex: F_1,43_=31, P<0.0001; effect of genotype: F_1,43_=0.1, P=0.8; subregion x sex interaction: F_1,43_=17, P=0.0002; all other interactions P>0.25). C) In the 25°C water maze, there were more Fos^+^ cells in females and in the dorsal granule cell layer, but there were no subregion-specific differences between the sexes (3 way RM ANOVA; effect of subregion: F_1,44_=74, P<0.0001; effect of sex: F_1,44_=4.2, P=0.048; effect of genotype: F_1,44_=1.9, P=0.2; subregion x sex interaction: F_1,44_=6, P=0.02; all other interactions P>0.09). D) In the 16°C water maze, there were more GAD67^+^Fos ^+^ cells females and in the ventral granule cell layer (mixed effects analysis; effect of subregion: F_1,42_=8, P=0.006; effect of sex: F_1,44_=8, P=0.009; effect of genotype: F_1,44_=0.0, P=0.99; all interactions: P>0.17. E) In the 25°C water maze, there were more GAD67^+^Fos^+^ cells in the ventral hippocampus but there were no sex or genotype differences (mixed effects analysis; effect of subregion: F_1,40_=5.2, P=0.03; effect of sex: F_1,43_=2.5, P=0.12; effect of genotype: F_1,43_=0.9, P=0.3; all interactions: P>0.24. Bars indicate mean ± s.e.m. mol, molecular layer; gcl, granule cell layer; hil, hilus; D, dorsal; V, ventral. *P<0.05, **P<0.01, ***P<0.001, ****P<0.0001.

### Immunohistochemistry

Animals were euthanized via overdose of isoflurane, and transcardially perfused with 4% paraformaldehyde in 0.1M phosphate buffer saline solution (PBS, pH 7.4). Brains were dissected and incubated in 4% paraformaldehyde for an additional 24 hours, after which they were placed in PBS with 0.1% sodium azide, and stored at 4°C. Prior to sectioning, brains were cryoprotected by incubation in 10% glycerol in PBS for 24 hours, followed by 20% glycerol for 48 hours. Brains were sectioned coronally through the hippocampus at 40 μm thickness using a freezing microtome and stored in cryoprotectant solution at −20°C until immunohistochemistry was completed.

For immunolabelling of doublecortin (DCX), one dorsal and one ventral section from each animal was mounted onto slides (Fisher, Superfrost) and left to dry for 24 hours. Slides were incubated in 0.1M citric acid and heated to an intermittent boil for 10 min for antigen-retrieval. Sections were then washed and incubated in PBS with 0.5% triton-X and 3% horse serum for 20 min. Tissue was then incubated in PBS with triton-X, with mouse-anti DCX monoclonal antibody (Santa Cruz Biotechnology, sc-271390, 1:100) at 4°C for 3 days. Sections were then rinsed in PBS, and incubated in biotinylated goat anti-mouse secondary antibody (Sigma, B0529,1:200) for 1 hour. Sections were washed, and treated with hydrogen peroxide (0.3%) in PBS for 30 min. Immunostaining was visualized through incubation in avidin-biotin-horseradish peroxidase (Vector Laboratories, Burlingame, CA) for 30 min, and subsequent treatment with cobalt-enhanced 3,3’-Diaminobenzidine chromogen (Sigma Fast Tablets, Sigma, St. Louis, MO). Sections were then counter-stained with cresyl-violet (0.1%), dehydrated, cleared with citrisolv (Thermofisher, Waltham, MA) and coverslipped with permount (Fisher).

For immunostaining of GFP, serial sections were incubated in mouse anti-GFP (DSHB, GFP-12E6, 1:100 in PBS with triton-X) for 24 hours, washed, incubated for 2 hours with donkey anti-mouse Alexa488 secondary antibody, washed, mounted onto slides and coverlipped with PVA-DABCO.

For immunostaining of c-Fos, sections were incubated in goat anti-c-Fos primary antibody (1:2000, Santa Cruz sc-52-G) in PBS-TX with horse serum for 3 days at 4°C. Sections were then washed 3 times in PBS-TX, and then incubated in secondary biotinylated donkey anti-goat antibody (1:250, Jackson Immunoresearch: 705-065-147) for 1 hour in PBS-TX with horse serum. The sections were then washed 3 times in PBS-TX, incubated in blocking solution (0.5%, Perkin Elmer: FP1020) for 30 min, before application of Streptavidin-HRP (1:100, NEL750) for 1 hour. Sections were then washed (3 x 5min) in PBS-TX, and incubated in Rhodamine (1:2000, Fisher Scientific: PI-46406) in PBS-TX and H_2_O_2_ (1:20,000) for 1 hour. Sections were then washed (3 x 5min) in PBS-TX, blocked for 30min in PBS-TX with horse serum, and then incubated in mouse anti-GAD67 primary antibody (1:1000, Millipore MAB5406) in PBS-TX with horse serum for 3 days at 4°C. Following GAD67 antibody incubation, sections were then washed 3 times in PBS-TX, and incubated in donkey anti-mouse Alexa 647 antibody (1:250, Invitrogen A-31571) for 1 hour. Tissue was then washed in PBS-TX (3 x 5min), and incubated in DAPI (1:1000) for 10 min. Lastly, sections were washed for (3 x 5min) in PBS, mounted onto glass slides, and coverslipped using PVA-Dabco mounting medium.

### Quantification of immunolabelling

Quantification of all immunolabelling was completed by an experimenter blind to the experimental conditions. For DCX, the number of immuno-positive cells was counted within the granule cell layer of the DG, using an Olympus CX41 bright-field microscope with a 40x objective. The number of immuno-positive cells were counted from 1 section of the septal/dorsal hippocampus (Bregma, −2.92 to −4.0 mm). Counts of DCX cells were also obtained from hippocampal sections which contained temporal/ventral hippocampus, although counts were not separated between the supra- and infra-pyramidal blades (Bregma, −5.76 to −6.2 mm). Intermediate and ventral DG was delineated at 4.5 mm relative to the interaural line. All counts of DCX positive cells were converted into densities based on the volume of the DG subregions.

For quantification of Fos immunoreactivity, a confocal microscope (Leica, SP8) was used to obtain representative z-stacks (40x objective), through the entire infrapyramidal and suprapyramidal blades of the DG, the medial and lateral blades of the ventral DG, and dorsal and ventral CA3. For each animal, an entire dorsal and ventral section was analyzed. Cells were counted as Fos-positive when the intensity of immunolabelling was more than twice that of neighboring, non-nuclei-containing, tissue in the hilus. To determine the percentage of GAD cells that also expressed Fos, Gad immunopositive cells were examined throughout the entire DG-CA3 and the proportion that expressed Fos at twice background levels was quantified.

Analyses of dendritic spine density were performed from z-stack images acquired with a 63x glycerol-immersion objective (NA 1.3). Images were 1024×1024 pixels in size, taken at 5x zoom, a speed of 400 Hz, and a z-height of 0.5 μm. For each neuron, images were acquired from the outer molecular layer (where lateral perforant path axons terminate), middle molecular layer (where medial perforant path axons terminate), and inner molecular layer (where mossy cell / commissural fibers terminate). All protrusions were counted as spines and mushroom spines were defined as having a head diameter ≥ 0.6 μm. A total of 14-37 cells per group, distributed equally across 3-5 animals/group, were analyzed.

Analyses of mossy fiber terminals were performed from z-stack images acquired with a 40x oil-immersion objective (NA 1.3). Images were 1024×1024 pixels in size, taken at 2x zoom, a speed of 400 Hz, and a z-height of 0.5 μm. The area of the large mossy terminal was measured from maximum intensity projections and the number of terminal-associated filopodia, more than 1μm in length, was also quantified as a proxy for GABAergic interneuron innervation (55, 56). Large mossy terminals and filopodia were categorized according to their position along the proximodistal CA3 axis, where CA3a is the curved distal portion of CA3, CA3c is proximal and enclosed within the blades of the DG, and CA3b is the intermediate CA3 region. A total of 59-122 large mossy terminals per group, distributed equally across 3-5 animals/group, were analyzed.

### Statistical Analysis

Analysis of water maze acquisition performance was performed using mixed-design repeated measures ANOVA with sex and genotype as between-subject factors and day of training as a within subject factor. The distribution of search strategies in WT and TK rats was analyzed by a Chi squared test with Bonferroni correction for multiple comparisons. Probe trial performance was analyzed with between-subject ANOVAs (sex x genotype). For behavioral experiments, 16°C and 25°C groups were typically analyzed and presented separately; in some cases we directly compared 16°C and 25°C groups to explore temperature effects. Cell densities were analyzed by mixed design repeated measures ANOVA with sex and genotype as between subject factors and dorsoventral subregion as a within subjects factor. Neuronal morphology (spines, boutons, filopodia) was analyzed by ANOVA with sex and treatment as between-subjects factors. For all ANOVAs, where significant interactions were detected, post-hoc comparisons were analyzed with Holm-Sidak tests. The significance level, *α*, was set at p=0.05 for all tests. In most cases, statistical results are presented in the figure legends alongside their respective data; for data that is not presented in figures, statistical results are presented in the results text.

### Sex and neurogenesis literature summary

To assess the degree to which sex has been included as a variable in studies of adult neurogenesis (Fig. 1), a Pubmed search was performed using the search terms “neurogenesis” and “dentate gyrus”. Results were binned into 5-year increments from 2001 to 2020 and 76-112 studies/bin (mean=98, distributed equally over the 5 years of a bin) were examined for whether they studied male, female or both male and female subjects, whether they formally analyzed their data by sex, or whether they did not report the sex of their subjects. To similarly assess inclusion of males and females in studies that have attempted to specifically manipulate neurogenesis and test behavioral consequences (“functional studies”), additional search terms were included (‘irradiation’, ‘tk’, ‘tmz’, ‘mam’, ‘arac’, ‘tamoxifen’, ‘chemogenetics’, and ‘optogenetics’; 97 results published between 2001 and 2021).

## RESULTS

### Inhibition of neurogenesis in male and female TK rats

To establish that neurogenesis was effectively inhibited along the dorsoventral axis of the DG in both male and female TK rats, we quantified the density of cells expressing the immature neuronal marker, doublecortin (DCX). As expected, in WT rats DCX^+^ cells were observed at the border of the granule cell layer and the hilus, in the subgranular zone (Fig. 2A). DCX^+^ cell density was dramatically reduced in both male and female TK rats, to less than 15% of levels found in WT littermates, consistent with recent work (48, 53, 57). This reduction was observed in the dorsal and ventral hippocampus, and there were no sex differences in the extent of neurogenesis reduction (Fig. 2B).

### In cold water, ablation of neurogenesis impairs spatial learning in male rats and improves spatial learning in female rats

Ablating neurogenesis typically does not impair learning a single spatial location in the water maze (2, 6, 7, 17–22). Since adult-born neurons regulate unconditioned responses to stressors (3, 23, 32), we hypothesized that stress or aversiveness may also reveal a role for new neurons in spatial learning. We therefore tested WT and TK rats in the spatial water maze at standard temperatures (25°C) or colder, more aversive temperatures (16°C).

In standard 25°C water, WT and TK rats learned to escape from the pool with similar latencies (Fig. 3A-C) and, in the probe trial, WT and TK rats displayed equivalent memory (Fig. 3D-F). We also observed sex differences in performance, where males escaped faster and spent more time in the target zone than females. We explored whether estrous stages influenced probe trial performance (but not after training trials, to avoid lavage impacts on subsequent behavior). The distribution of WT and TK rats across the 4 stages of the estrous cycle did not differ (**χ**^2^=1.3, P=0.7) but rats in proestrus displayed better memory than rats that were not in proestrus, an effect that was comparable for both WT and TK rats (Fig. 3G).

At 16°C, blocking neurogenesis altered learning in both males and females, but in opposite directions: male TK rats located the platform faster but female TK rats located it faster, compared to their WT counterparts (Fig. 4A-C). As in 25°C water, WT male rats located the platform faster than WT females. These effects could not be explained by differences in swim speed (Supplementary Fig. 1). A similar pattern was observed when we analyzed ideal path error, a measure of the cumulative positional error relative to the platform that is not influenced by differences in swim speed or path length (52): at 16°C female TK rats had a lower path error and male TK rats had a greater path error, relative to WT controls (Supplementary Fig. 2).

On the probe trial, TK rats spent less time searching in the target zone (Fig. 4D-F). This pattern was stronger in male TK rats but the ANOVA interaction (sex x time spent in target zone) was not significant. Following the 16°C probe trial, the estrous distribution of female WT and TK rats did not differ (**χ**^2^=2.7, P=0.4) and there was no effect of estrous stage on probe trial performance (Fig. 4G).

To rule out the possibility that behavioral differences were due to nonspecific physiological effects caused by cold water, we measured body temperature in a separate group of rats. At both 16°C and 25°C, body temperature was lowest after day 1 training, was lower on day 1 in females than in males, but not different between WT and TK rats (Supplementary Fig. 3). Male TK rats weighed slightly less than male WT rats, consistent with previous studies showing that neurogenesis inhibition can sometimes reduce weight (19, 48) (8%; WT: 480 ± 8g, TK: 441 ± 8g; mean ± s.e.m.). However, female WT and TK rats were not different (3%; WT: 279 ± 5g, TK: 270 ± 6g; 2 way ANOVA; effect of genotype: F_1,116_=11, P=0.001; genotype x sex interaction: F_1,116_=4, P=0.049; female WT vs TK: P=0.6; male WT vs TK: P=0.0001). Furthermore, neither body weight nor body temperature correlated with learning and memory performance at 16°C or 25°C, suggesting that water temperature did not differentially impact sexes or genotypes due to hypothermic effects (Supplementary Tables 1 & 2). Finally, to rule out the possibility that TK impairments and enhancements in learning are due to nonspecific effects of the GFAP-TK transgene, we trained additional WT and TK rats that did not receive valganciclovir treatment. Here, no genotype differences were observed at 16°C or 25°C water temperatures (Supplementary Fig. 4).

To gain insight into navigational strategies employed during learning, we analyzed search patterns with Pathfinder software (52). Generally, rats displayed increasing use of spatially-specific search strategies over days of testing (Fig. 5). Specifically, they shifted from thigmotaxic and random searches, or searches that covered multiple areas of the pool equally, to searches that were biased towards the escape platform with increasing precision. Male TK rats relied less on spatially-specific search strategies than their WT counterparts. Consistent with their faster escape latency, female TK rats tended to display more spatially-specific searches than their WT counterparts but this difference was not statistically significant. Consistent with the latency and path error data, search strategies did not differ between WT and TK rats tested at 25°C (Supplementary Fig. 5).

Behavioral sex differences often reflect differences in strategy (58, 59). We therefore explored whether maze aversiveness caused males and females to employ different navigational strategies in the water maze. Female WT rats responded strongly to cold temperature, and spent less time searching randomly and at the edge of the pool, and more time performing spatial searches in the center of the pool and near the platform. Temperature-dependent changes in search strategy were absent in female TK that lacked neurogenesis (Supplementary Fig. 6). In contrast, male WT rats employed similar strategies at both 16°C and 25°C, but blocking neurogenesis led to temperature-dependent differences, where TK males performed fewer spatially precise searches in 16°C water. Thus, neurogenesis promotes aversiveness-related changes in search strategy in females but it promotes consistent search strategies in males.

### Blocking neurogenesis did not alter the HPA response

Neurogenesis regulates the HPA axis in mice (23) and cold temperatures can enhance water maze learning via glucocorticoid-dependent mechanisms (49). We therefore explored whether neurogenesis regulates HPA axis function in rats at baseline and after learning. Consistent with previous work in mice (23), we found no neurogenesis-related changes in baseline circadian HPA function. Corticosterone levels were highest at the onset of darkness, they were higher in females, but they did not differ between WT and TK rats (Supplementary Fig. 7). When corticosterone was measured 30 min after the first day of acquisition training, both WT and TK rats displayed high levels of corticosterone, which did not differ between genotypes. Corticosterone levels also did not differ between rats trained at 16°C vs 25°C. When normalized to escape latency, i.e. time spent in the water, there was a tendency for greater corticosterone levels at 16°C but this did not reach statistical significance. A subset of rats that were subjected to the full 4 days of testing displayed HPA habituation, but no corticosterone differences were observed between genotypes or temperatures. Thus, females elicit a stronger HPA response than males, but neurogenesis-associated behavioral differences at 16°C are not due to differences in HPA output.

### Activity-induced Fos expression is modulated by sex and dorsoventral location, but not neurogenesis

Behaviorally-relevant DG neuronal populations express the activity-dependent immediate-early gene, c-Fos (60–62). To determine whether blocking neurogenesis alters neuronal population activity in males and females, we quantified Fos expression in excitatory principal cell populations in DG-CA3, in both WT and TK rats (Fig. 6). Notably, Fos activation was never different between WT and TK rats. However, more dentate granule neurons were active in females than in males, particularly at 16°C (74% more at 16°C, 24% more at 25°C). There were also strong dorsoventral gradients of activity: at 16°C, females had ~2x greater Fos levels in the dorsal DG compared to the ventral DG or the dorsal DG of males. In contrast, males trained at 16°C did not display a significant dorsoventral gradient of activity. At 25°C, females also displayed a strong dorsoventral gradient of activity but in males this effect was weaker with only TK rats having significantly greater Fos activation in the dorsal DG. To explore whether Fos levels differed across training temperatures, we pooled genotypes and performed a sex x temperature ANOVA (dorsal and ventral subregions combined). A significant interaction revealed that females had more Fos^+^ cells when trained at 16°C than at 25°C; males did not differ (effect of sex: F_1,91_=30, P<0.0001, effect of temperature: F_1,91_=3.3, P=0.07; interaction F_1,91_=6.8, P=0.01; female 16°C vs 25°C: P=0.008; male 16°C vs 25°C: P=0.95). Since a shift in reliance on the ventral-to-dorsal hippocampus mediates the progression toward spatially-specific search strategies (63), we explored relationships between Fos activation of dorsal vs ventral hippocampus with performance on the acquisition and retrieval stages of testing, however, no significant correlations were observed (data not shown).

Since adult-born neurons can influence DG-CA3 activity via efferent connections with inhibitory interneurons (56, 64), we quantified Fos^+^ inhibitory, GAD67-expressing neurons in DG-CA3 (Fig. 6D,E). In rats trained at 16°C, there was a strong dorsoventral gradient of activity in GAD67^+^ cells, with greater activity in the ventral DG than in the dorsal DG. There was also significantly greater activation of GAD67^+^ cells in females than in males, but no differences due to loss of adult neurogenesis. Sparse activation precluded a robust subregional analysis but, when analyzed by DG-CA3 subregion, sex differences in GAD67^+^ cell activation were observed in the molecular layer, granule cell layer and hilus+CA3 (female > male), but the dorsoventral difference was specific to the granule cell layer (ventral > dorsal; Supplementary Fig. 8A-C). At 25°C, fewer GAD67^+^ cells were activated (mixed effects analysis; effect of temperature: F_1,91_=8.2, P=0.005) and the dorsoventral gradient (V > D) was weaker. In contrast to rats trained at 16°C, there were no sex differences in activation of GAD67^+^ cells in rats trained at 25°C. Finally, at 25°C there also were no differences between genotypes. A similar pattern was observed in the granule cell layer and hilus+CA3 region (Supplementary Fig. 8D-F).

### Training- and sex-dependent morphological plasticity in adult-born neurons

Functionally-relevant morphological features of adult-born neurons develop during the weeks and months post-mitosis (65–67), and can be modified by spatial learning (68, 69). To examine sex differences in experience-dependent plasticity we labelled adult-born neurons with retrovirus and analyzed GFP^+^ spines and presynaptic terminals as morphological proxies for afferent and efferent connectivity (Fig. 7). At baseline, in naïve home cage rats, there were no differences in spine density between adult-born neurons from male and female rats. However, in male rats, training at 16°C specifically elevated spine density compared to rats that were untrained or trained at 25°C, and compared to female rats trained at 16°C. This effect was observed throughout the molecular layer. In both males and females, regardless of treatment, spine density increased with distance from the cell soma as previously observed (65) (Supplementary Fig. 10). The density of large, mushroom spines was not altered by training (Fig. 7C).

**Figure 7:**
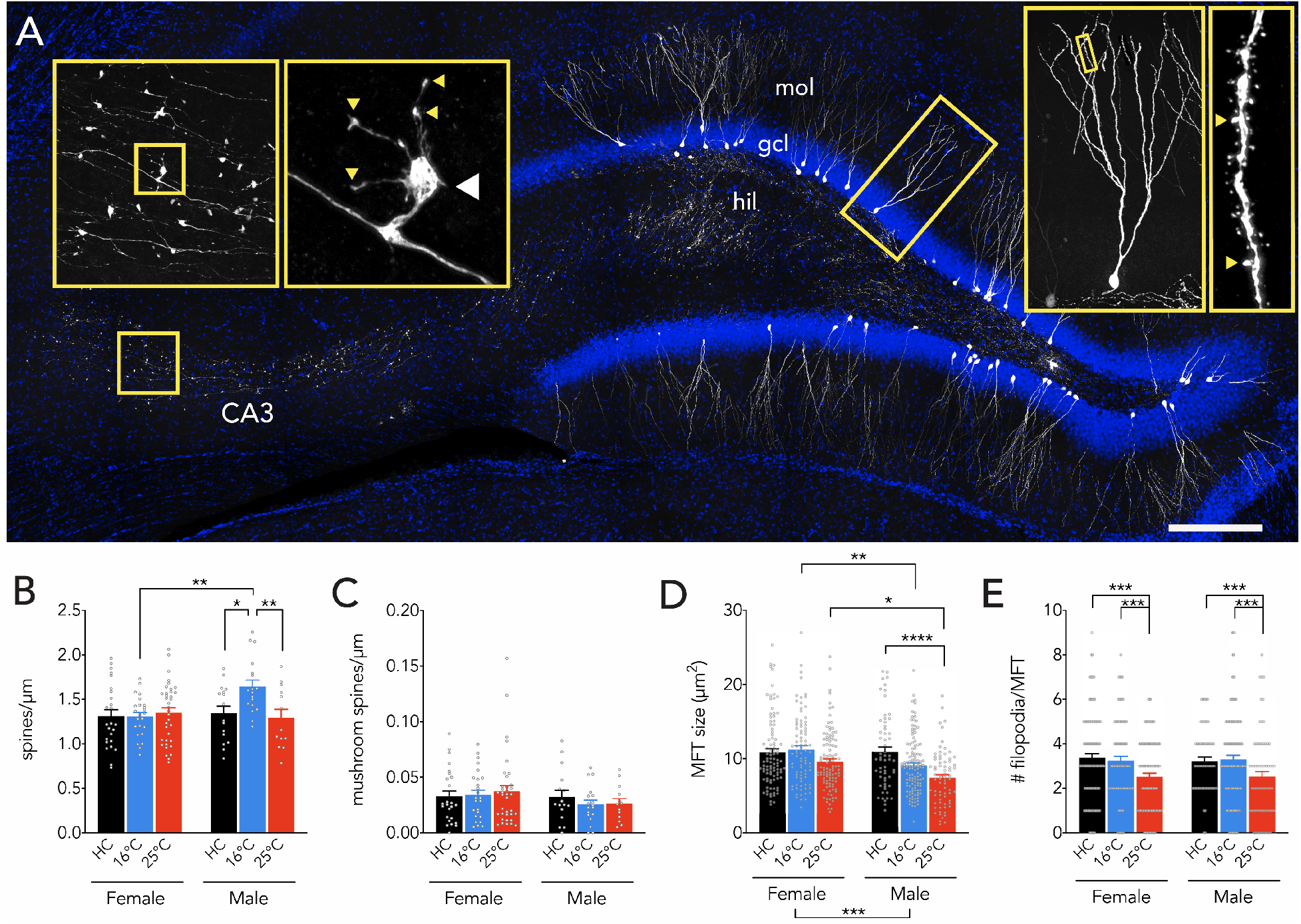
Training reveals sex- and temperature-dependent plasticity. A) Retroviral GFP labelling of adult-born neurons in the dentate gyrus, with axons projecting to CA3. Right insets display an isolated neuron (reconstructed across sections, hence the greater number of dendrites) and dendrite (arrowheads indicate mushroom spines). Left insets display a large mossy fiber terminal (MFT); white arrowhead indicates the MFT and yellow arrowheads indicate putative presynaptic filopodial contacts onto inhibitory interneurons. Scale bar, 250 μm. hil, hilus; gcl, granule cell layer; mol, molecular layer. B) Adult-born neuron spine density was selectively increased in male rats that were trained at 16°C (2 way anova; effect of treatment: F_2,127_=3.1, P=0.0495; effect of sex: F_1,127_=3.2, P=0.08; interaction: F_2,127_=4.2, P=0.02; Male home cage (HC) vs 16°C: P=0.01; male HC vs 25°C: P=0.7; male 16°C vs 25°C: P=0.009; female group comparisons all P>0.95; male vs female at 16°C: P=0.003; male vs female HC and male vs female at 25°C both P>0.8). C) Adult-born neuron mushroom spine density was not altered by sex or training (2 way anova; effect of treatment: F_2,127_=0.1, P=0.12; effect of sex: F_1,127_=2.0, P=0.15; interaction: F_2,127_=0.5, P=0.6;). D) MFTs were larger in adult-born neurons from female rats, an effect that was driven by greater training-related reduction in terminal size in males (2 way anova; effect of sex, F_1,539_ = 14, P=0.0002; effect of training condition F_2,539_=13, P<0.0001; interaction, F_2,539_= 3.5, P=0.03; male HC vs male 16°C, P=0.06; male HC vs male 25°C, P<0.0001; female HC vs female 16°C and 25°C both P>0.18; male HC vs female HC, P=0.9; male 16°C vs female 16°C, P=0.005; male 25°C vs female 25°C, P=0.01). E) The number of MFT-associated filopodia, putative synapses onto inhibitory neurons, was reduced in the 25°C group but was not different between sexes (2 way anova; effect of training condition, F_2,545_=9, P<0.0001, effect of sex, F_1,545_=0, P=0.9; interaction, F_2,545_=0.1, P=0.9). Bars indicate mean ± s.e.m. mol, molecular layer; gcl, granule cell layer; hil, hilus. *P<0.05, **P<0.01, ***P<0.001, ****P<0.0001.

Finally, we examined the large mossy fiber terminals that excite CA3 pyramidal neurons. No sex differences were observed between naïve, home cage control rats. However, only in males, training decreased mossy fiber terminal size, an effect that was greatest in the 25°C group (Fig. 7D). In both females and males, 25°C training also reduced the number of filopodial extensions that protrude off of mossy fiber boutons, putative synapses onto inhibitory interneurons (Fig. 7E).

## DISCUSSION

Sex modulates hippocampal memory, plasticity and physiology (70). And while there is also evidence that sex regulates the addition and activation of new neurons (71), relatively few studies have formally investigated sex and none have identified sex differences in the behavioral consequences of manipulating adult neurogenesis (Fig. 1). Here we report that blocking neurogenesis caused female rats to learn faster and male rats to learn slower, relative to intact rats in a spatial water maze at aversive 16°C temperatures. These findings were not confounded by genotype differences in swim speed, body weight or body temperature, and they were not present in TK rats that were not treated with valganciclovir (and therefore had intact neurogenesis). Whereas new neurons were morphologically equivalent at baseline, learning evoked distinct patterns of pre- and post-synaptic plasticity depending on sex. Our study therefore provides new evidence that adult-born neurons make unique sex-dependent contributions to spatial learning under stress and have distinct plasticity profiles in male and female rats.

### Temperature-dependent spatial functions of newborn neurons

While some have reported acquisition and short term reference memory deficits in the spatial water maze in neurogenesis-deficient animals (14, 69, 72), a majority of studies have found intact spatial learning (2, 6, 7, 15, 17–22), raising questions about the necessity of adult neurogenesis for spatial learning. Our findings indicate that the degree of stress and/or aversiveness present at the time of learning is critical (as suggested by (73)). Indeed, there is ample evidence that neurogenesis regulates innate fear and anxiety-like behaviors in response to stressful and aversive stimuli (23–25, 27–29). And while stress is known to potently modulate hippocampal memory, few studies have examined a role for neurogenesis in learning as a function of stress: one study found that neurogenesis is critical for context fear memory when mice receive a single, but not multiple, footshocks (33); another found that TK rats made more errors in a dry spatial maze only when an aversive odor was present (32). That we found no learning differences at 25°C suggests that neurogenesis may be particularly important for spatial learning under conditions of higher stress. Notably, we did not find differences in HPA activation between rats trained at 16°C and 25°C. However, 16°C training did lead to greater hippocampal recruitment (in females), greater dorsoventral differences in hippocampal activation, differences in strategy usage, and it caused a greater reduction in body temperature. Thus, 16°C water evoked physiological changes and behaviors that are consistent with the concept of a stressor as a threatening stimulus that perturbs an organism from baseline, necessitating an adaptive or homeostatic response (74).

### Sex differences in the behavioral function of adult-born neurons

We found that blocking neurogenesis led to opposite behavioral outcomes in females and males which, to our knowledge, is the first report of sex differences in the behavioral function of neurogenesis. To date, sex differences in function have gone undetected because few studies have compared male and female animals that have altered neurogenesis (“functional” studies). In our attempt to comprehensively survey the literature (Fig. 1), we counted only 4 functional studies that have reported data by sex or included sex as a variable in their statistical analyses (9, 57, 75, 76).

It is typically understood that neurogenesis benefits cognition and so it may seem paradoxical that blocking neurogenesis improved water maze learning in females. However, it has been repeatedly demonstrated that males and females can display opposite patterns of hippocampal-dependent learning, with manipulations facilitating performance in males in some paradigms and facilitating performance in females in others (42–44). Our findings also may seem paradoxical if it is assumed that “faster is better” in the water maze. It is increasingly well-documented that sex differences in learning tasks can reflect strategy differences rather than frank differences in learning ability (58, 77) and escape latencies cannot reveal differences in strategy and navigational choice that may be highly adaptive (15). Here we found that male rats that lacked neurogenesis performed more general searches, but female neurogenesis-deficient rats tended to (nonsignificantly) perform more spatially specific searches. While it is common to view spatially-specific searches as “better”, generalized search has clear advantages in cases where a spatial goal moves to a new or unexpected location (78, 79). Thus, one possibility is that neurogenesis adjusts search/memory specificity differently, increasing it in males and perhaps decreasing it in females. That females trained at 16°C had significantly higher levels of Fos in the dorsal DG indicates that there are clear sex differences in regional hippocampal recruitment, which could impact the adoption of precise search strategies (63).

Another possibility, related to the fact that neurogenesis effects were selectively observed in 16°C water, is that emotional functions of neurogenesis were differentially engaged by stress. In other studies, stress impairs spatial learning in males and is either without effect, or actually improves learning, in females (42, 43). These divergent effects may reflect differential effects of stress on cognition (males) and hyperarousal (females) (80). Since neurogenesis ablation mimics some features of the stressed brain (e.g. structural atrophy) (81, 82), possibly male learning was impaired by dysregulated integration of stress and learning, and females learned faster due to heightened arousal and attention effects. A role for attentional processes is also suggested by recent work showing that blocking neurogenesis reduces orienting responses to distractor stimuli (83), an effect that may explain why TK rats are faster to navigate a dry spatial maze in the presence of an aversive, but irrelevant, mint odor (32). Given sex differences in processing object arrays and configurations (70), blocking neurogenesis may differentially alter water maze cue processing such that females are less susceptible to distraction from irrelevant cues (leading to faster escape) but males are less attentive to relevant cues (leading to slower escape).

Finally, insights into the potential adaptive significance of neurogenesis also come from our analyses across temperatures (Supplementary Fig. 6). Intact females were highly sensitive to temperature: 16°C shifted females away from random and wall-focused search, toward the center of the pool and the specific area of the platform. In contrast, TK females were not different at 16°C and 25°C. Thus, in females, neurogenesis promotes changes in strategy according to the aversiveness of the situation. In males, neurogenesis promoted equivalent strategy usage 16°C and 25°C, which could also be adaptive in cases where performance needs to remain stable despite perturbations from external forces.

### Sex differences in hippocampal subregional activation

To investigate possible subregional and cellular mechanisms we examined activity-dependent Fos expression along the dorsoventral axis in male and female rats that did, or did not, have adult neurogenesis. While previous studies have reported that ablating neurogenesis can increase (13, 27, 64) or decrease (53, 84) activity in the hippocampus, here we found no effect on global activity amongst dentate granule cells. Newborn neurons also target inhibitory interneurons (56, 64), whose activity regulates the precision of hippocampal-dependent memory (85, 86). However, we also observed no changes in inhibitory recruitment in TK rats relative to WT rats. While these findings suggest that neurogenesis ablation did not affect behavior by altering hippocampal activity, it is possible that activity differences were present early in training, when behavioral sex differences were more prominent.

Little is known about how dorsoventral subregions of the hippocampus are activated in males and females by training in the standard spatial water maze. Here, we found that females consistently had greater levels of DG activity than males, particularly at 16°C. This was largely driven by elevated Fos levels in the dorsal hippocampus, a finding that builds on previous evidence that the spatial water maze recruits dorsal more than ventral DG (60). However, whereas that study only included males, here we find that the dorsoventral gradient is significantly stronger in females. Elevated Fos in dorsal vs ventral DG was mirrored by an opposite gradient of Fos in GAD67^+^ inhibitory cells, suggesting that regional activity is controlled by local inhibitory circuits. Since the temporal progression of water maze learning strategies involves sequential recruitment of ventral to dorsal hippocampus (63) we explored relationships between water maze performance (latency, path error, strategy specificity on acquisition and probe trials) and activity in the dorsal and ventral DG. However, we found no consistent correlations, suggesting that other forms of activity and plasticity may be more tightly linked to performance.

### Sex differences in morphological plasticity of adult-born neurons

To our knowledge, this is the first study to examine functionally-relevant morphological features of adult-born neurons in males and females. At baseline, we observed no differences in spine density or mossy fiber terminal size between the sexes. However, water maze training induced plasticity of excitatory synaptic structures but only in males. Since blocking neurogenesis impaired 16°C learning in males, 16°C-induced spinogenesis may be important for learning under stress in males, possibly allowing for greater association of sensory information from entorhinal cortical inputs. Somewhat surprisingly, training reduced the size of mossy fiber terminals in males. These findings are reminiscent of early work showing that the CA3 pyramidal neuron apical dendrites, which are targeted by mossy fiber axons, undergo greater stress-induced plasticity in males than in females (87). Given the link between mossy fiber terminal size and synaptic strength (88, 89), training likely reduced synaptic strength in male rats trained at 25°C, suggesting that new neurons in males may play a weaker role in memory under less aversive conditions. Likewise, we observed fewer filopodial protrusions in both males and females trained at 25°C, suggesting that new neurons are less likely to recruit inhibitory circuits in less aversive conditions, an effect that could reduce memory precision (85, 86).

## ACKNOWLEDGEMENTS

This work was supported by CIHR operating funds (Catalyst Grant: Sex as a variable in biomedical research) to JSS, a Michael Smith Foundation for Health Research Scholar Award to JSS, a CIHR New Investigator Award to JSS, and a Michael Smith Foundation for Health Research Postdoctoral Fellowship to TPO. The authors thank Stephanie Hipkin for assistance with the Figure 1 literature review.

## REFERENCES

1. T. J. Shors, et al., Neurogenesis in the adult is involved in the formation of trace memories. Nature 410, 372–376 (2001).

2. T. J. Shors, D. A. Townsend, M. Zhao, Y. Kozorovitskiy, E. Gould, Neurogenesis may relate to some but not all types of hippocampal-dependent learning. Hippocampus 12, 578–584 (2002).

3. D. o Seo, M. A. Carillo, S. C.-H. Lim, K. F. Tanaka, M. R. Drew, Adult Hippocampal Neurogenesis Modulates Fear Learning through Associative and Nonassociative Mechanisms. The Journal of neuroscience: the official journal of the Society for Neuroscience 35, 11330–11345 (2015).

4. C. D. Clelland, et al., A functional role for adult hippocampal neurogenesis in spatial pattern separation. 325, 210–213 (2009).

5. G. Winocur, J. M. Wojtowicz, M. Sekeres, J. S. Snyder, S. Wang, Inhibition of neurogenesis interferes with hippocampus-dependent memory function. Hippocampus 16, 296–304 (2006).

6. M. D. Saxe, et al., Ablation of hippocampal neurogenesis impairs contextual fear conditioning and synaptic plasticity in the dentate gyrus. Proceedings of the National Academy of Sciences of the United States of America 103, 17501–17506 (2006).

7. S. Jessberger, et al., Dentate gyrus-specific knockdown of adult neurogenesis impairs spatial and object recognition memory in adult rats. Learning & memory (Cold Spring Harbor, NY) 16, 147–154 (2009).

8. C. A. Denny, N. S. Burghardt, D. M. Schachter, R. Hen, M. R. Drew, 4-to 6-week-old adult-born hippocampal neurons influence novelty-evoked exploration and contextual fear conditioning. Hippocampus 22, 1188–1201 (2012).

9. E. C. Cope, et al., Adult-Born Neurons in the Hippocampus Are Essential for Social Memory Maintenance. Eneuro 7, ENEURO.0182-20.2020 (2020).

10. T. Kitamura, et al., Adult neurogenesis modulates the hippocampus-dependent period of associative fear memory. Cell 139, 814–827 (2009).

11. D. Kumar, et al., Sparse Activity of Hippocampal Adult-Born Neurons during REM Sleep Is Necessary for Memory Consolidation. Neuron (2020) https:/doi.org/10.1016/j.neuron.2020.05.008.

12. K. G. Akers, et al., Hippocampal Neurogenesis Regulates Forgetting During Adulthood and Infancy. Science (New York, NY) 344, 598–602 (2014).

13. N. S. Burghardt, E. H. Park, R. Hen, A. A. Fenton, Adult-born hippocampal neurons promote cognitive flexibility in mice. Hippocampus 22, 1795–1808 (2012).

14. A. Garthe, J. Behr, G. Kempermann, Adult-generated hippocampal neurons allow the flexible use of spatially precise learning strategies. PLoS ONE 4, e5464 (2009).

15. R. Q. Yu, M. Cooke, D. R. Seib, J. Zhao, J. S. Snyder, Adult neurogenesis promotes efficient, nonspecific search strategies in a spatial alternation water maze task. Behav Brain Res 376, 112151 (2019).

16. A. A. Swan, et al., Characterization of the role of adult neurogenesis in touch-screen discrimination learning. Hippocampus 24, 1581–1591 (2014).

17. T. M. Madsen, P. E. G. Kristjansen, T. G. Bolwig, G. Wörtwein, Arrested neuronal proliferation and impaired hippocampal function following fractionated brain irradiation in the adult rat. Neuroscience 119, 635–642 (2003).

18. J. Raber, et al., Radiation-induced cognitive impairments are associated with changes in indicators of hippocampal neurogenesis. Radiation research 162, 39–47 (2004).

19. J. S. Snyder, N. S. Hong, R. J. McDonald, J. M. Wojtowicz, A role for adult neurogenesis in spatial long-term memory. Neuroscience 130, 843–852 (2005).

20. C. A. Blaiss, et al., Temporally Specified Genetic Ablation of Neurogenesis Impairs Cognitive Recovery after Traumatic Brain Injury. J Neurosci 31, 4906–4916 (2011).

21. C. G. Nickell, K. R. Thompson, J. R. Pauly, K. Nixon, Recovery of Hippocampal-Dependent Learning Despite Blunting Reactive Adult Neurogenesis After Alcohol Dependence. Adv Neurol 6, 83–101 (2020).

22. J. O. Groves, et al., Ablating Adult Neurogenesis in the Rat Has No Effect on Spatial Processing: Evidence from a Novel Pharmacogenetic Model. PLoS genetics 9, e1003718 (2013).

23. J. S. Snyder, A. Soumier, M. Brewer, J. Pickel, H. A. Cameron, Adult hippocampal neurogenesis buffers stress responses and depressive behaviour. Nature 476, 458–461 (2011).

24. J.-M. Revest, et al., Adult hippocampal neurogenesis is involved in anxiety-related behaviors. Molecular psychiatry 14, 959–967 (2009).

25. M. L. Lehmann, R. A. Brachman, K. Martinowich, R. J. Schloesser, M. Herkenham, Glucocorticoids Orchestrate Divergent Effects on Mood through Adult Neurogenesis. The Journal of neuroscience: the official journal of the Society for Neuroscience 33, 2961–2972 (2013).

26. A. Surget, et al., Antidepressants recruit new neurons to improve stress response regulation. Molecular psychiatry 16, 1177–1188 (2011).

27. C. Anacker, et al., Hippocampal neurogenesis confers stress resilience by inhibiting the ventral dentate gyrus. Nature 559, 1–22 (2018).

28. D. C. Lagace, et al., Adult hippocampal neurogenesis is functionally important for stress-induced social avoidance. Proceedings of the National Academy of Sciences 107, 4436–4441 (2010).

29. T. J. Schoenfeld, et al., New neurons restore structural and behavioral abnormalities in a rat model of PTSD. Hippocampus 29, 848–861 (2019).

30. B. Roozendaal, J. L. McGaugh, Memory modulation. Behavioral neuroscience 125, 797–824 (2011).

31. D. A. Bangasser, T. J. Shors, Critical brain circuits at the intersection between stress and learning. Neuroscience and biobehavioral reviews 34, 1223–1233 (2010).

32. T. J. Schoenfeld, J. A. Smith, A. N. Sonti, H. A. Cameron, Adult neurogenesis alters response to an aversive distractor in a labyrinth maze without affecting spatial learning or memory. Hippocampus (2020) https:/doi.org/10.1002/hipo.23267.

33. M. R. Drew, C. A. Denny, R. Hen, Arrest of adult hippocampal neurogenesis in mice impairs single-but not multiple-trial contextual fear conditioning. Behavioral neuroscience 124, 446–454 (2010).

34. R. C. Kessler, M. Petukhova, N. A. Sampson, A. M. Zaslavsky, H. Wittchen, Twelve-month and lifetime prevalence and lifetime morbid risk of anxiety and mood disorders in the United States. Int J Method Psych 21, 169–184 (2012).

35. C. Chow, J. R. Epp, S. E. Lieblich, C. K. Barha, L. A. M. Galea, Sex differences in neurogenesis and activation of new neurons in response to spatial learning and memory. Psychoneuroendocrinology 38, 1236–1250 (2013).

36. S. Yagi, et al., Sex Differences in Maturation and Attrition of Adult Neurogenesis in the Hippocampus. Eneuro 7, ENEURO.0468-19.2020 (2020).

37. S. Yagi, C. Chow, S. E. Lieblich, L. A. M. Galea, Sex and strategy use matters for pattern separation, adult neurogenesis, and immediate early gene expression in the hippocampus. Hippocampus 26, 87–101 (2016).

38. J. M. Juraska, J. M. Fitch, C. Henderson, N. Rivers, Sex differences in the dendritic branching of dentate granule cells following differential experience. Brain research 333, 73–80 (1985).

39. S. G. Warren, A. G. Humphreys, J. M. Juraska, W. T. Greenough, LTP varies across the estrous cycle: enhanced synaptic plasticity in proestrus rats. Brain research 703, 26–30 (1995).

40. T. J. Shors, C. Chua, J. Falduto, Sex differences and opposite effects of stress on dendritic spine density in the male versus female hippocampus. The Journal of neuroscience: the official journal of the Society for Neuroscience 21, 6292–6297 (2001).

41. H. E. Scharfman, N. J. MacLusky, Differential regulation of BDNF, synaptic plasticity and sprouting in the hippocampal mossy fiber pathway of male and female rats. Neuropharmacology 76 Pt C, 696–708 (2014).

42. V. Luine, Sex differences in chronic stress effects on memory in rats. Stress (Amsterdam, Netherlands) 5, 205–216 (2002).

43. C. D. Conrad, et al., Acute stress impairs spatial memory in male but not female rats: influence of estrous cycle. Pharmacology, biochemistry, and behavior 78, 569–579 (2004).

44. D. A. Bangasser, T. J. Shors, The hippocampus is necessary for enhancements and impairments of learning following stress. Nature neuroscience 10, 1401–1403 (2007).

45. A. K. Beery, I. Zucker, Sex bias in neuroscience and biomedical research. Neuroscience and biobehavioral reviews 35, 565–572 (2011).

46. K. A. Huckleberry, R. M. Shansky, The unique plasticity of hippocampal adult-born neurons: Contributing to a heterogeneous dentate. Hippocampus (2021) https:/doi.org/10.1002/hipo.23318.

47. K. Roughton, M. Kalm, K. Blomgren, Sex-dependent differences in behavior and hippocampal neurogenesis after irradiation to the young mouse brain. The European journal of neuroscience 36, 2763–2772 (2012).

48. J. S. Snyder, et al., A Transgenic Rat for Specifically Inhibiting Adult Neurogenesis. eneuro 3 (2016).

49. C. Sandi, M. Loscertales, C. Guaza, Experience-dependent facilitating effect of corticosterone on spatial memory formation in the water maze. The European journal of neuroscience 9, 637–642 (1997).

50. B. Salehi, M. I. Cordero, C. Sandi, Learning under stress: the inverted-U-shape function revisited. Learning & memory (Cold Spring Harbor, NY) 17, 522–530 (2010).

51. M. Gallagher, R. Burwell, M. Burchinal, Severity of spatial learning impairment in aging: development of a learning index for performance in the Morris water maze. Behavioral neuroscience 107, 618–626 (1993).

52. M. B. Cooke, et al., Pathfinder: open source software for analyzing spatial navigation search strategies. F1000research 8, 1521 (2019).

53. D. R. Seib, et al., Hippocampal neurogenesis promotes preference for future rewards. Mol Psychiatr, 1–19 (2021).

54. A. C. McLean, N. Valenzuela, S. Fai, S. A. L. Bennett, Performing vaginal lavage, crystal violet staining, and vaginal cytological evaluation for mouse estrous cycle staging identification. Journal of visualized experiments: JoVE, e4389 (2012).

55. L. Acsády, A. Kamondi, A. Sík, T. Freund, G. Buzsáki, GABAergic cells are the major postsynaptic targets of mossy fibers in the rat hippocampus. The Journal of neuroscience: the official journal of the Society for Neuroscience 18, 3386–3403 (1998).

56. L. Restivo, Y. Niibori, V. Mercaldo, S. A. Josselyn, P. W. Frankland, Development of Adult-Generated Cell Connectivity with Excitatory and Inhibitory Cell Populations in the Hippocampus. The Journal of neuroscience: the official journal of the Society for Neuroscience 35, 10600–10612 (2015).

57. D. R. Seib, E. Chahley, O. Princz-Lebel, J. S. Snyder, Intact memory for local and distal cues in male and female rats that lack adult neurogenesis. PLoS ONE 13, e0197869–15 (2018).

58. R. M. Shansky, Sex differences in behavioral strategies: avoiding interpretational pitfalls. Curr Opin Neurobiol 49, 95–98 (2018).

59. W. G. Brake, J. M. Lacasse, Sex differences in spatial navigation: the role of gonadal hormones. Curr Opin Behav Sci 23, 176–182 (2018).

60. J. S. Snyder, R. Radik, J. M. Wojtowicz, H. A. Cameron, Anatomical gradients of adult neurogenesis and activity: young neurons in the ventral dentate gyrus are activated by water maze training. Hippocampus 19, 360–370 (2009).

61. X. Liu, et al., Optogenetic stimulation of a hippocampal engram activates fear memory recall. Nature 484, 381–385 (2012).

62. S. R. Erwin, et al., A Sparse, Spatially Biased Subtype of Mature Granule Cell Dominates Recruitment in Hippocampal-Associated Behaviors. Cell Reports 31, 107551 (2020).

63. S. Ruediger, D. Spirig, F. Donato, P. Caroni, Goal-oriented searching mediated by ventral hippocampus early in trial-and-error learning. Nature neuroscience 15, 1563–1571 (2012).

64. L. J. Drew, et al., Activation of local inhibitory circuits in the dentate gyrus by adult-born neurons. Hippocampus (2015) https:/doi.org/10.1002/hipo.22557.

65. J. D. Cole, et al., Adult-born hippocampal neurons undergo extended development and are morphologically distinct from neonatally-born neurons Prolonged development of adult-born neurons. J Neurosci Official J Soc Neurosci, JN-RM-1665-19 (2020).

66. J. T. Gonçalves, et al., In vivo imaging of dendritic pruning in dentate granule cells. Nature neuroscience (2016) https:/doi.org/10.1038/nn.4301.

67. C. Zhao, E. M. Teng, R. G. Summers, G.-L. Ming, F. H. Gage, Distinct Morphological Stages of Dentate Granule Neuron Maturation in the Adult Mouse Hippocampus. The Journal of neuroscience: the official journal of the Society for Neuroscience 26, 3–11 (2006).

68. S. Tronel, et al., Spatial learning sculpts the dendritic arbor of adult-born hippocampal neurons. Proceedings of the National Academy of Sciences 107, 7963–7968 (2010).

69. V. Lemaire, et al., Long-lasting plasticity of hippocampal adult-born neurons. The Journal of neuroscience: the official journal of the Society for Neuroscience 32, 3101–3108 (2012).

70. W. A. Koss, K. M. Frick, Sex differences in hippocampal function. Journal of neuroscience research 95, 539–562 (2017).

71. S. Yagi, L. A. M. Galea, Sex differences in hippocampal cognition and neurogenesis. Neuropsychopharmacol 44, 200–213 (2019).

72. D. Dupret, et al., Spatial relational memory requires hippocampal adult neurogenesis. PLoS ONE 3, e1959 (2008).

73. A. Dranovsky, E. D. Leonardo, Is there a role for young hippocampal neurons in adaptation to stress? Behavioural brain research 227, 371–375 (2012).

74. Y. M. Ulrich-Lai, J. P. Herman, Neural regulation of endocrine and autonomic stress responses. Nature reviews Neuroscience 10, 397–409 (2009).

75. K. A. Huckleberry, et al., Dorsal and ventral hippocampal adult-born neurons contribute to context fear memory. Neuropsychopharmacology: official publication of the American College of Neuropsychopharmacology 43, 2487–2496 (2018).

76. L. N. Miller, C. Weiss, J. F. Disterhoft, Genetic Ablation of Neural Progenitor Cells Impairs Acquisition of Trace Eyeblink Conditioning. eneuro 6, ENEURO.0251-19.2019 (2019).

77. N. C. Tronson, Focus on females: a less biased approach for studying strategies and mechanisms of memory. Curr Opin Behav Sci 23, 92–97 (2018).

78. B. A. Richards, et al., Patterns across multiple memories are identified over time. Nature neuroscience 17, 981–986 (2014).

79. R. J. Steele, R. G. Morris, Delay-dependent impairment of a matching-to-place task with chronic and intrahippocampal infusion of the NMDA-antagonist D-AP5. Hippocampus 9, 118–136 (1999).

80. D. A. Bangasser, S. R. Eck, A. M. Telenson, M. Salvatore, Sex differences in stress regulation of arousal and cognition. Physiol Behav 187, 42–50 (2018).

81. T. J. Schoenfeld, H. C. McCausland, H. D. Morris, V. Padmanaban, H. A. Cameron, Stress and Loss of Adult Neurogenesis Differentially Reduce Hippocampal Volume. Biological Psychiatry 82, 1–34 (2017).

82. R. J. Schloesser, et al., Atrophy of pyramidal neurons and increased stress-induced glutamate levels in CA3 following chronic suppression of adult neurogenesis. Brain structure & function (2013) https:/doi.org/10.1007/s00429-013-0532-8.

83. C. S. S. Weeden, J. C. Mercurio, H. A. Cameron, A role for hippocampal adult neurogenesis in shifting attention toward novel stimuli. Behav Brain Res 376, 112152 (2019).

84. L. R. Glover, T. J. Schoenfeld, R.-M. Karlsson, D. M. Bannerman, H. A. Cameron, Ongoing neurogenesis in the adult dentate gyrus mediates behavioral responses to ambiguous threat cues. PLoS biology 15, e2001154 (2017).

85. S. Ruediger, et al., Learning-related feedforward inhibitory connectivity growth required for memory precision. Nature 473, 514–518 (2011).

86. N. Guo, et al., Dentate granule cell recruitment of feedforward inhibition governs engram maintenance and remote memory generalization. Nature Publishing Group 24, 438–449 (2018).

87. L. A. Galea, et al., Sex differences in dendritic atrophy of CA3 pyramidal neurons in response to chronic restraint stress. Neuroscience 81, 689–697 (1997).

88. I. Galimberti, et al., Long-Term Rearrangements of Hippocampal Mossy Fiber Terminal Connectivity in the Adult Regulated by Experience. Neuron 50, 749–763 (2006).

89. I. Galimberti, E. Bednarek, F. Donato, P. Caroni, EphA4 signaling in juveniles establishes topographic specificity of structural plasticity in the hippocampus. Neuron 65, 627–642 (2010).

