## Supplementary Data for "Inhibiting adult neurogenesis differentially affects spatial learning in females and males"

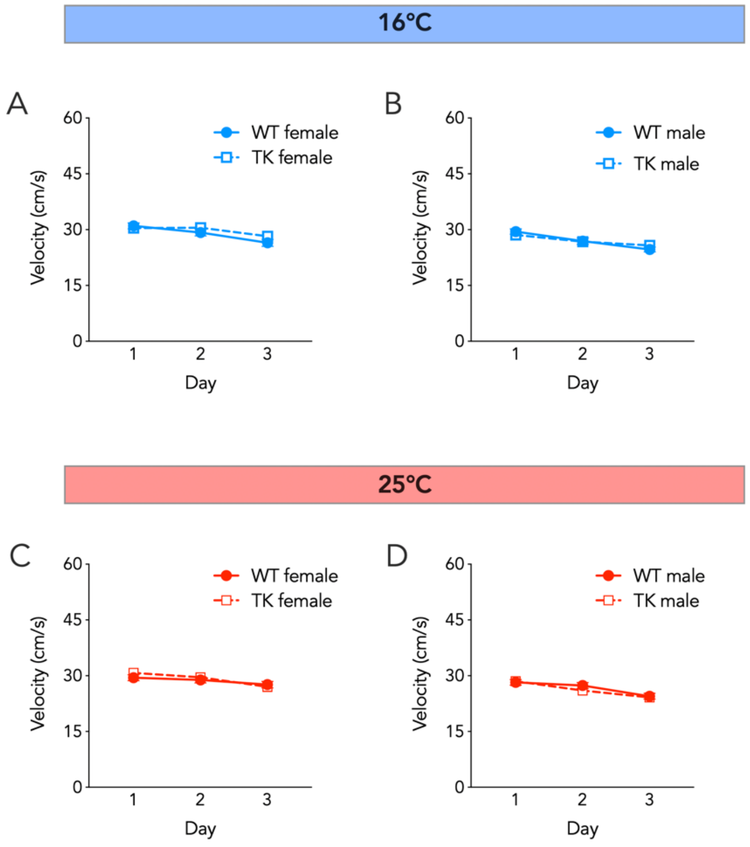


**Supplementary Figure 1: Water maze swim speed**. A,B) In the 16°C water maze, females swam faster than males, swim speed declined over days, and there were no differences between WT and TK rats (3 way anova; effect of sex: $F_{1,78} = 8$, P=0.006; effect of day: $F_{2,156} = 32$, P<0.0001; effect of genotype: $F_{1,78} = 0.1$, P=0.7; all interactions P>0.05). C,D) In the 25°C water maze, females swam faster than males, swim speed declined over days, and there were no differences between WT and TK rats (3 way anova; effect of sex: $F_{1,76} = 13$, P=0.0005; effect of day: $F_{2,152} = 35$, P<0.0001; effect of genotype: $F_{1,76} = 0$, P=1; all interactions P>0.25). Symbols reflect mean ± standard error. *P<0.05, ****P<0.0001. N=18-22 per group.

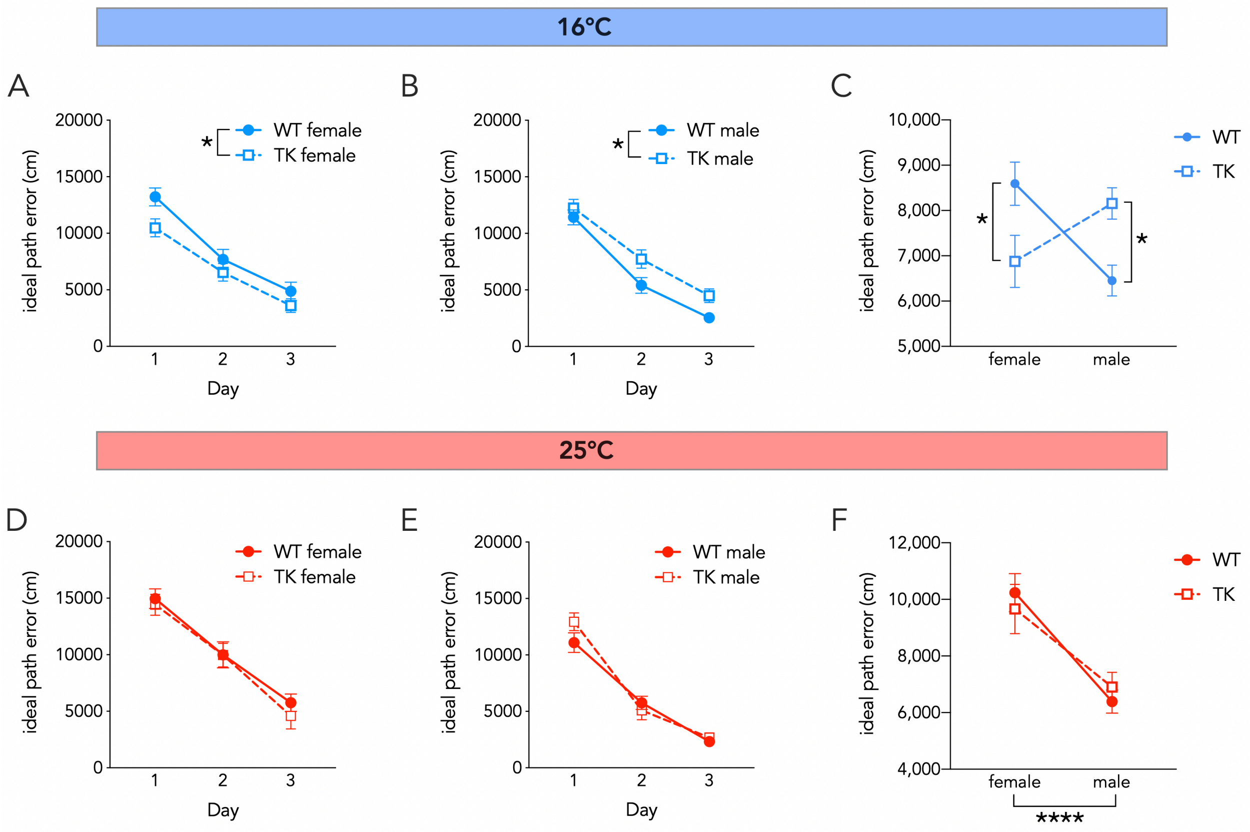


**Supplementary Figure 2: Analyses of ideal path error in WT and TK rats.** A,B) In the 16°C water maze, ideal path error decreased over days (3 way anova; effect of genotype, $F_{1,78}$ = 0.0, P=0.99; effect of day, $F_{2,156}$ = 133, P<0.0001; effect of sex, $F_{1,78}$ = 1.0, P=0.3). Blocking neurogenesis increased ideal path error in males but decreased it in females (genotype x sex interaction: $F_{1,78}$ = 16, P=0.0002; male WT vs male TK, P = 0.03; female WT vs female TK, P = 0.03). Male WT rats also had lower ideal path error score than female WT rats (P=0.005). C) Average ideal path error scores during training. D,E) In the 25°C water maze, ideal path error was not different between genotypes, but decreased over days and was lower for males than females (3 way anova; effect of genotype, $F_{1,76}$ = 0.0, P=0.96; effect of day, $F_{2,152}$ = 202, P<0.0001; effect of sex, $F_{1,76}$ = 28, P<0.0001; all interactions, P>0.05). F) Average trial ideal path error scores during training. Symbols reflect mean ± standard error. *P<0.05, ****P<0.0001. N = 18-22 per group.

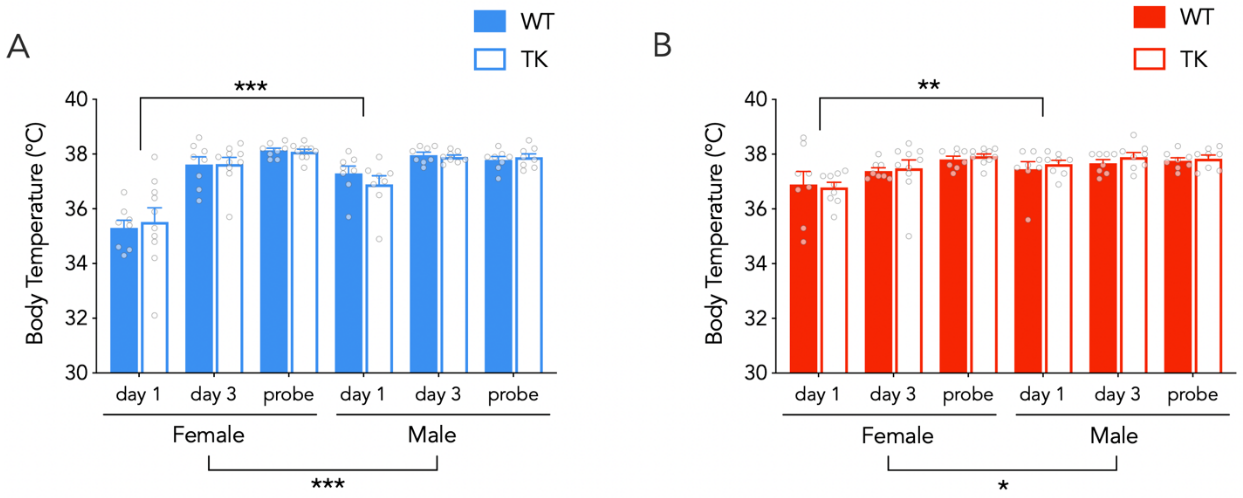


**Supplementary Figure 3: Water maze body temperatures**. A) At 16°C, post-testing body temperatures were lowest on day 1, were lower on day 1 in females than in males, but were not different between WT and TK rats (3 way repeated measures ANOVA; effect of day: $F_{2,60}$=52, P<0.0001; effect of sex: $F_{1,30}$=14.1, P=0.0008; effect of genotype: $F_{1,30}$=0.04, P=0.8; day x sex interaction: $F_{2,64}$=16, P<0.0001; all genotype interactions P>0.5; day 1 vs day 3 and probe both P<0.0001; day 1 female vs male: P<0.001; days 3 and probe female vs male: P=0.4). B) At 25°C, post-testing body temperatures were lowest on day 1, were lower on day 1 in females than in males, but were not different between WT and TK rats (3 way repeated measures mixed effects model; effect of day: $F_{2,59}$=11, P<0.0001; effect of sex: $F_{1,30}$=4.8, P=0.04; effect of genotype: $F_{1,30}$=0.4, P=0.5; all genotype interactions P>0.6; day x sex interaction: $F_{2,63}$=4.2, P=0.02; day 1 vs day 3, P=0.005, day 1 vs probe, P<0.0001; day 1 female vs male: P=0.005, day 3 and probe female vs male: P=0.7). Bars reflect mean ± standard error. *P<0.05, **P<0.01, ***P<0.001.

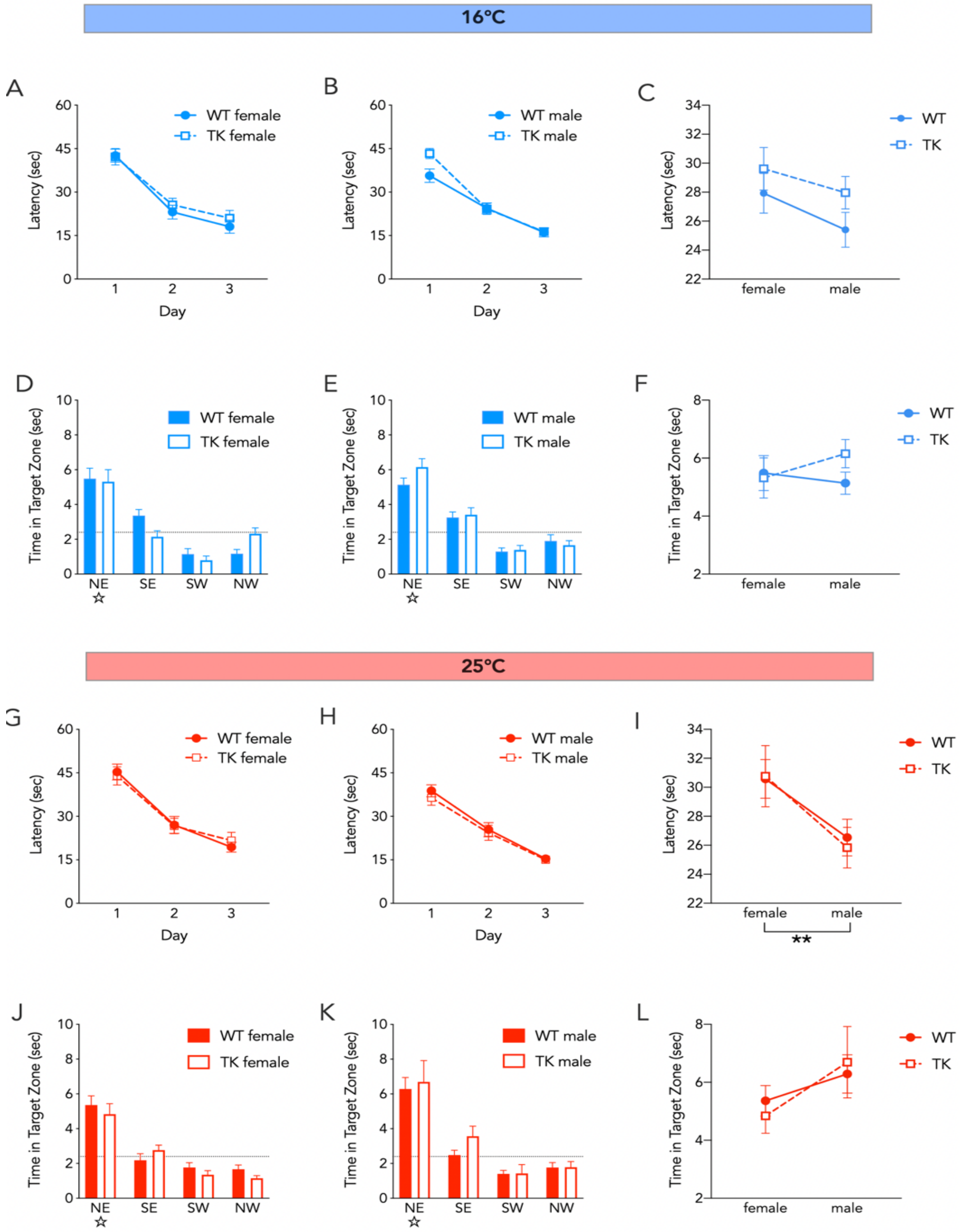


**Supplementary Figure 4: Water maze performance in WT and TK rats that were not treated with valganciclovir**. A-C) Spatial water maze learning at 16°C was similar in WT and TK rats (3-way anova; effect of day, $F_{2,156} = 130$, $P<0.0001$; effect of sex, $F_{1,78} = 2.6$, $P=0.1$; effect of genotype, $F_{1,78} = 2.7$, $P=0.1$; interactions all $P>0.1$). D-F) 16°C probe trial performance was similar in WT and TK rats (2-way anova; effect of genotype, $F_{1,78} = 0.6$, $P=0.4$; effect of sex, $F_{1,78} = 0.2$, $P=0.6$; interaction, $F_{1,78} = 1.2$, $P=0.3$). G-I) Spatial water maze learning at 25°C was similar in WT and TK rats (3-way anova; effect of day, $F_{2,154} = 103$, $P<0.0001$; effect of sex, $F_{1,77} = 10$, ** =0.0028; effect of genotype, $F_{1,77} = 0.1$, $P=0.7$; interactions all $P>0.29$). J-L) 25°C probe trial performance was similar in WT and TK rats (2-way anova; effect of genotype, $F_{1,77} = 0$, $P=0.9$; effect of sex, $F_{1,77} = 3.4$, $P=0.06$; interaction, $F_{1,77} = 0.3$, $P=0.5$). Bars and symbols indicate mean ± standard error.

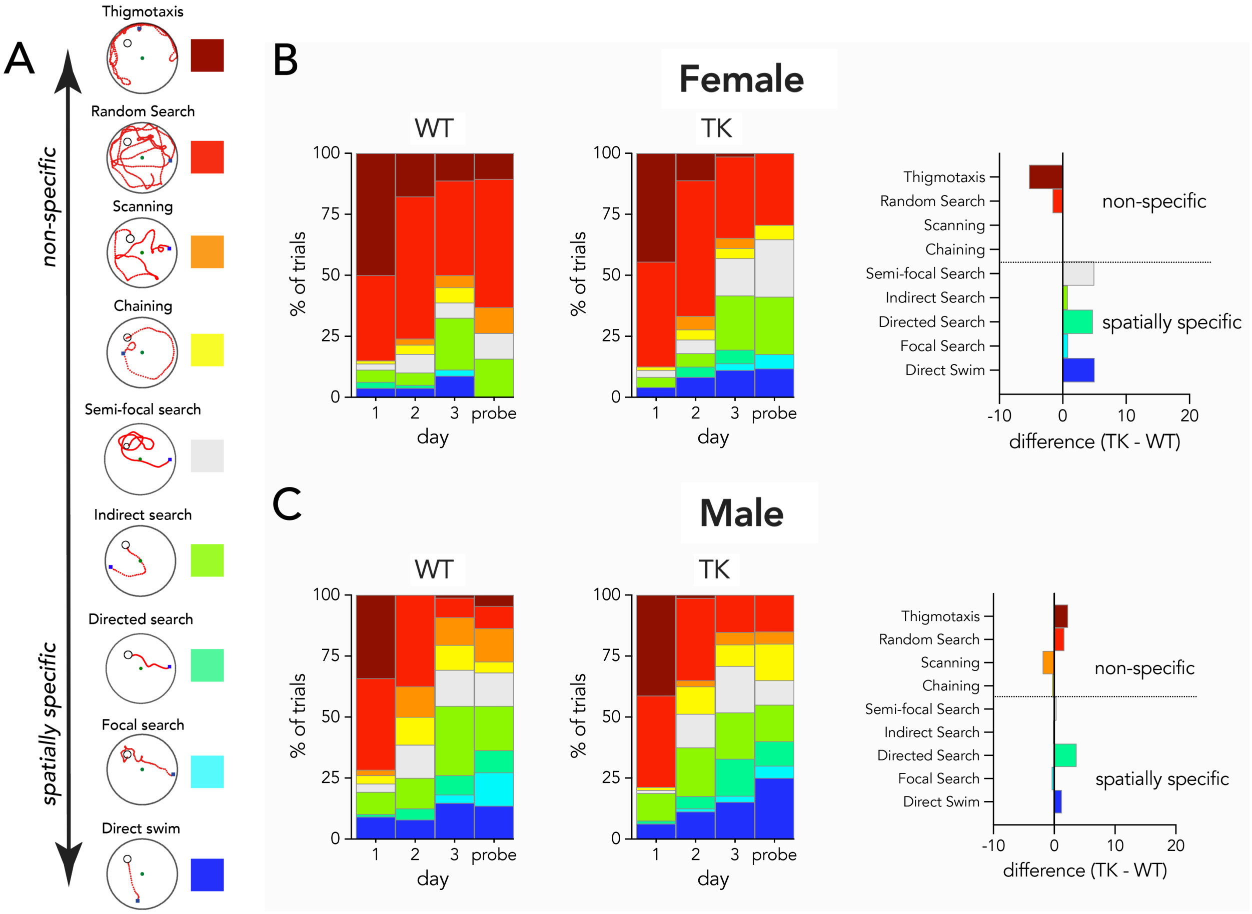


**Supplementary Figure 5: Similar strategies used by WT and TK rats at 25°C.** A) Example trials illustrating various search strategies classified by Pathfinder, organized by degree of spatial specificity relative to the target. B) Strategies employed by female WT (left) and TK (middle) rats and the relative change in TK rats (right). The distribution of strategies in female TK rats was not significantly different from female WT rats ($\chi^2$ = 9, P = 0.3). C) Strategies employed by male WT (left) and TK (middle) rats and the weighted difference between TK and WT rats. Reducing neurogenesis did not alter the distribution of strategies used by male rats ($\chi^2$ = 11, P = 0.4).

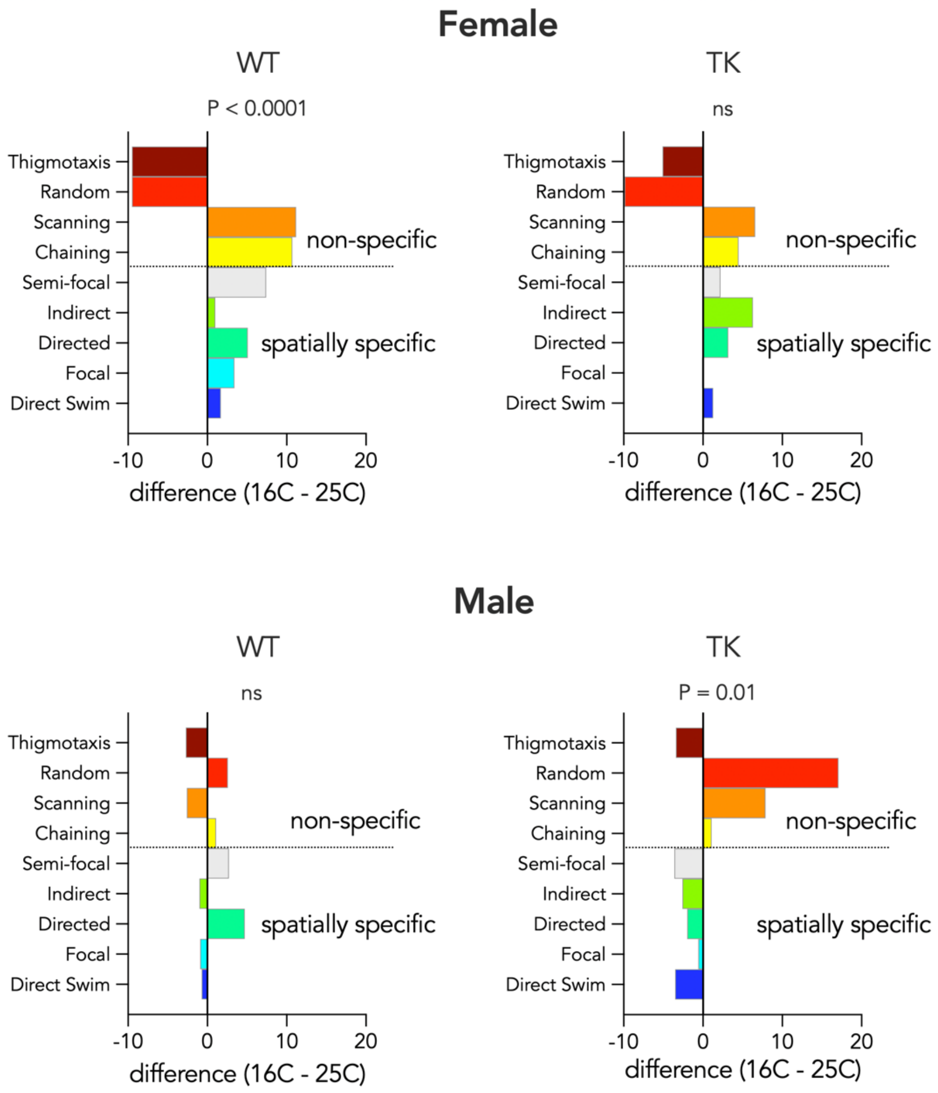


**Supplementary Figure 6: Temperature-related changes in search strategy**. Graphs show weighted strategy differences between rats trained at 16°C and 25°C. A) WT females employed different strategies as a function of temperature, and performed fewer thigmotactic and random searches at 16°C ($\chi^2 = 38$, $P < 0.0001$) B) TK females' strategy did not differ across temperatures ($\chi^2 = 18$, $P = 0.02$). C) Male WT rats did not alter strategies as a function of temperature ($\chi^2 = 9.3$, $P = 0.3$). D) Male TK rats performed fewer spatially specific searches at 16°C ($\chi^2 = 23$, $P = 0.01$). Bonferroni-corrected $P = 0.0125$. ns, not significant.

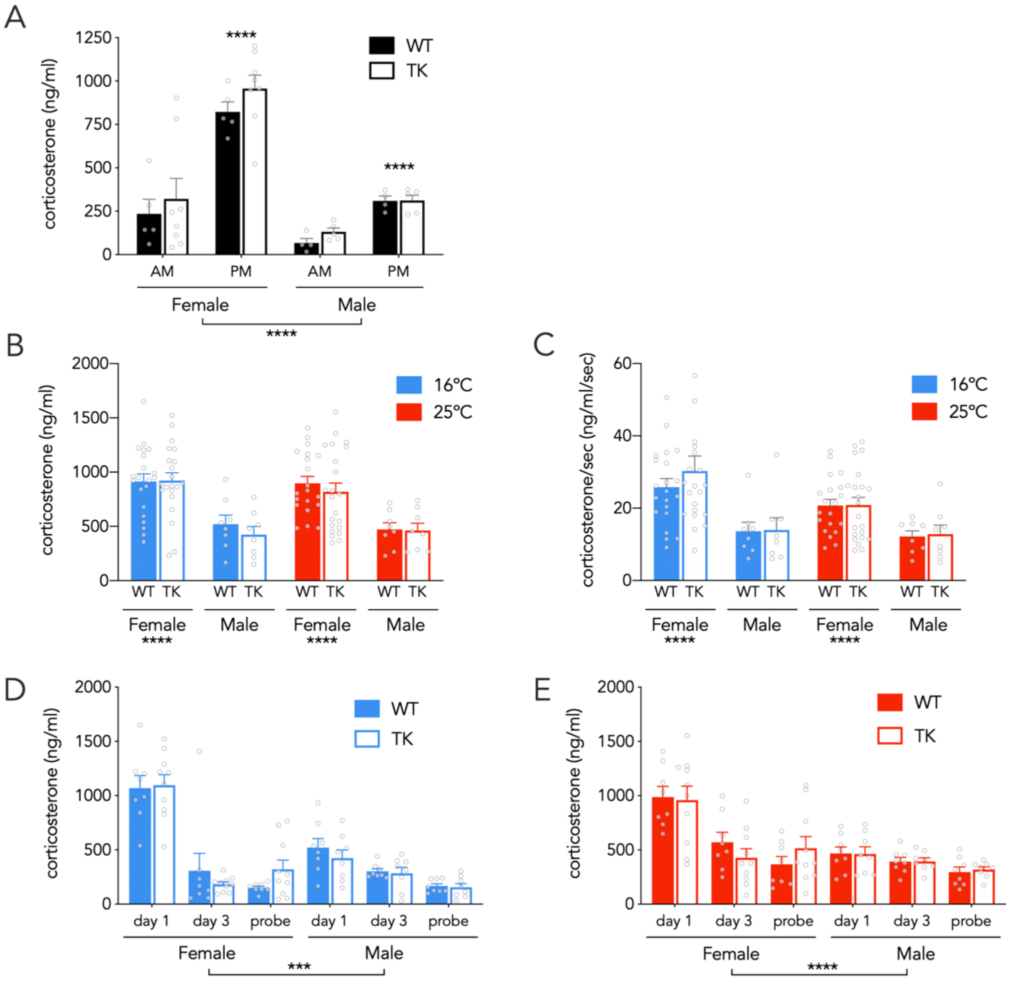


**Supplementary Figure 7: Similar HPA reactivity in WT and TK rats**. A) Baseline corticosterone levels are modulated by circadian phase and sex, but not neurogenesis (3 way ANOVA; effect of time of day: $F_{1,18}$=55, P<0.0001; effect of sex: $F_{1,18}$=34, P<0.0001; effect of genotype: $F_{1,18}$1.2, P=0.3; genotype interactions all P>0.5). B) On day 1 of acquisition, corticosterone levels were higher in females but were not different between genotypes or between rats trained at 16°C or 25°C (3 way ANOVA; effect of sex: $F_{1,108}$=48, P<0.0001; effect of genotype: $F_{1,108}$=0.5, P=0.5; effect of temperature: $F_{1,108}$=0.3, P=0.6; genotype interactions all P≥0.5). C) Day 1 corticosterone, normalized to time spent in the water maze, was higher in females but not significantly different between genotypes or rats trained at 16°C or 25°C (3 way ANOVA; effect of sex: $F_{1,108}$=22, P<0.0001; effect of genotype: $F_{1,108}$=0.3, P=0.6; effect of temperature: $F_{1,108}$=3.2, P=0.07; interactions all P>0.2). D) HPA activity habituated over days in the 16°C water maze but did not differ between genotypes (3 way ANOVA; effect of day: $F_{2,60}$=96, P<0.0001; effect of sex: $F_{1,30}$=13, P=0.001; effect of genotype: $F_{1,30}$=0.0, P=0.9; genotype interactions all P>0.2). E) HPA activity habituated over days in the 25°C water maze but did not differ between genotypes (3 way ANOVA; effect of day: $F_{2,60}$=26, P<0.0001; effect of sex: $F_{1,30}$=18, P=0.0002; effect of genotype: $F_{1,30}$=0.0, P=1; genotype interactions all P>0.3). Bars indicate mean ± standard error.

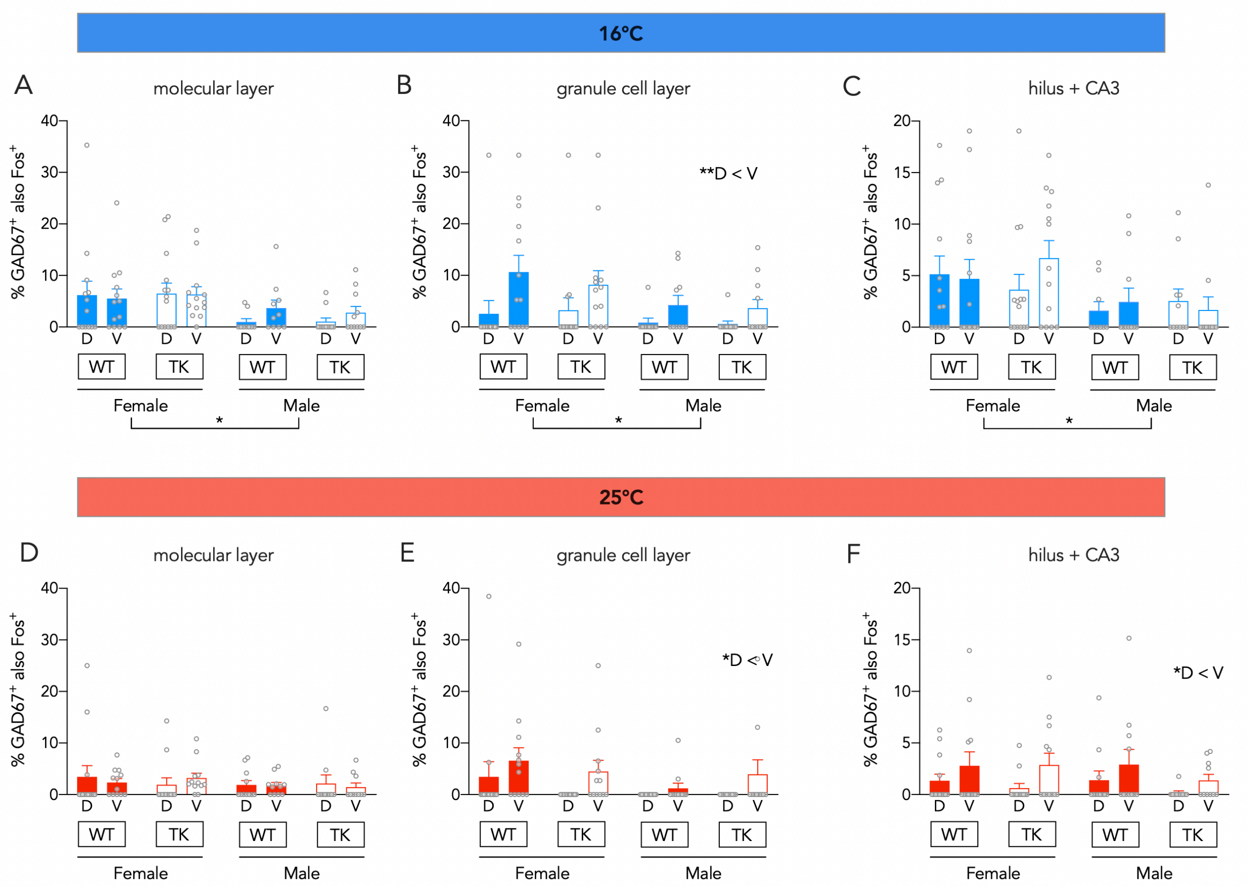


**Supplementary Figure 8: Activation of GAD67$^+$ interneurons across DG-CA3 subregions**. A) At 16°C, in the DG molecular layer, Fos expression was greater in GAD67$^+$ cells in females than in males (mixed effects analysis; effect of subregion: $F_{1,42}$=1.1, P=0.3; effect of sex: $F_{1,44}$=6.2, P=0.02; effect of genotype: $F_{1,44}$=0.0, P=0.97; all interactions: P>0.09. B) At 16°C, in the DG granule cell layer, Fos expression was greater in GAD67$^+$ cells in the ventral DG and in females (mixed effects analysis; effect of subregion: $F_{1,86}$=8.5, P=0.0046; effect of sex: $F_{1,86}$=5.2, P=0.02; effect of genotype: $F_{1,86}$=0.2, P=0.7; all interactions: P>0.3. C) At 16°C, in the combined hilar + CA3 region, Fos expression was greater in GAD67$^+$ cells in females than in males (mixed effects analysis; effect of subregion: $F_{1,42}$=0.5, P=0.5; effect of sex: $F_{1,44}$=5.6, P=0.02; effect of genotype: $F_{1,44}$=0.0, P=0.9; all interactions: P>0.16. D) At 25°C, in the DG molecular layer, Fos expression in GAD67$^+$ cells did not vary across sexes, subregions or genotypes (mixed effects analysis; effect of subregion: $F_{1,40}$=0.0, P=0.9; effect of sex: $F_{1,43}$=1.0, P=0.3; effect of genotype: $F_{1,43}$=0.0, P=0.9; all interactions: P>0.38. E) At 25°C, in the DG granule cell layer, Fos expression was greater in GAD67$^+$ cells in the ventral DG (mixed effects analysis; effect of subregion: $F_{1,83}$=5.3, P=0.02; effect of sex: $F_{1,83}$=2.8, P=0.1; effect of genotype: $F_{1,83}$=0.3, P=0.6; all interactions: P>0.14. F) At 25°C, in the hilus+CA3, Fos expression was greater in GAD67$^+$ cells in the ventral DG (mixed effects analysis; effect of subregion: $F_{1,40}$=7.3, P=0.01; effect of sex: $F_{1,43}$=0.3, P=0.6; effect of genotype: $F_{1,43}$=1.2, P=0.3; all interactions: P>0.46. Bars indicate mean ± s.e.m. D, dorsal; V, ventral. *P<0.05, **P<0.01

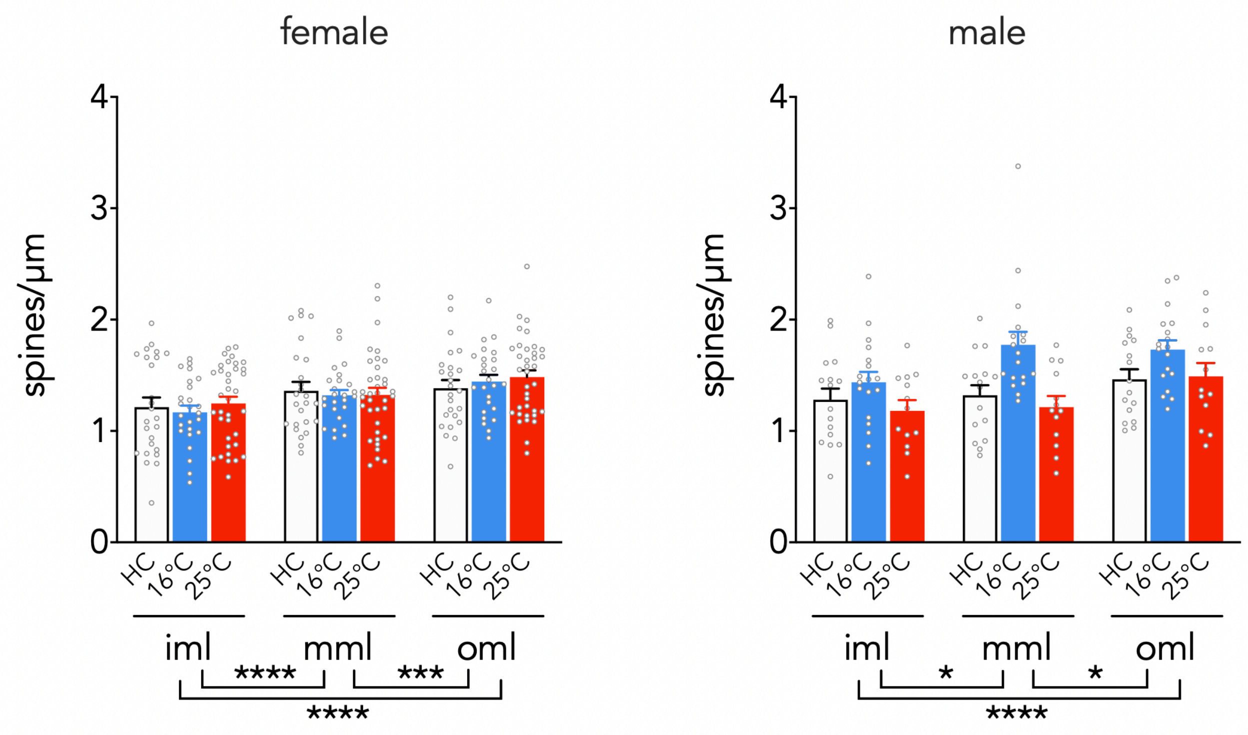


**Supplementary Figure 9: Molecular layer subregion spine densities**. Spine density increased across molecular layer subregions in both males and females (iml < mml < oml). Training did not differentially alter spine densities as a function of subregion. Males: 2 way repeated measure anova; effect of subregion, $F_{2,88}$=10.4, P<0.0001; treatment x subregion interaction: $F_{4,88}$=2.1, P=0.09. Females: effect of subregion, $F_{2,166}$=29, P<0.0001; treatment x subregion interaction: $F_{4,166}$=1.2, P=0.3. *P<0.05, ***P<0.001, ****P<0.0001. Bars reflect mean ± standard error.

| | Day 1 Latency | Day 2 Latency | Day 3 Latency | Probe |
|---|---|---|---|---|
| 16°C Males (N=36) | 0.13 (0.45) | -0.19 (0.26) | -0.21 (0.22) | 0.08 (0.63) |
| 16°C Females (N=24) | -0.05 (0.81) | -0.23 (0.28) | -0.13 (0.54) | 0.16 (0.45) |
| 25°C Males (N=36) | -0.04 (0.81) | 0.24 (0.16) | -0.24 (0.15) | -0.10 (0.56) |
| 25°C Females (N=24) | 0.23 (0.27) | 0.22 (0.28) | 0.06 (0.77) | -0.32 (0.15) |

**Supplementary Table 1: Correlations between body weight and learning and memory**. Pearson r correlation coefficients and respective P values in brackets. Body weight was not significantly correlated with performance on the water maze.

| | Day 1 Latency | Day 3 Latency | Probe |
|---|---|---|---|
| 16°C Males (N=16) | -0.32 (0.23) | -0.45 (0.08) | 0.06 (0.63) |
| 16°C Females (N=18) | -0.34 (0.17) | -0.30 (0.22) | 0.19 (0.44) |
| 25°C Males (N=16) | -0.07 (0.79) | -0.11 (0.70) | -0.14 (0.62) |
| 25°C Females (N=18) | -0.58 (0.02) | -0.48 (0.05) | 0.20 (0.44) |

**Supplementary Table 2: Correlations between body temperature and learning and memory**. Pearson r correlation coefficients and respective P values in brackets. Body temperature was not significantly correlated with performance on the water maze (where Bonferroni-corrected P = 0.0042)
